# Distinct roles of dentate gyrus and medial entorhinal cortex inputs for phase precession and temporal correlations in the hippocampal CA3 area

**DOI:** 10.1101/2022.06.07.491859

**Authors:** Siavash Ahmadi, Takuya Sasaki, Marta Sabariego, Christian Leibold, Stefan Leutgeb, Jill K. Leutgeb

## Abstract

The hippocampal CA3 subregion is a densely connected recurrent circuit that supports memory consolidation and retrieval by generating and storing sequential neuronal activity patterns that reflect recent experience. While theta phase precession is thought to be critical for generating sequential activity during memory encoding, the circuit mechanisms that support this computation across hippocampal subregions are unknown. By analyzing CA3 network activity in the absence of each of its theta modulated excitatory inputs, we show necessary and unique contributions of the dentate gyrus (DG) and the medial entorhinal cortex (MEC) to phase precession. DG inputs are essential for generating the preferential spiking of CA3 cells during late theta phases and for organizing the temporal order of neuronal firing, while MEC inputs modulate the general precision of phase precession. A computational model that accounts for the empirical findings suggests that DG inputs affect the phase and MEC inputs affect the amplitude of inhibitory subnetworks. Our results thus identify a novel and unique functional role of the DG for the generation of sequence coding in the CA3 recurrent circuit.

## Introduction

Although it is well established that the hippocampus supports episodic and spatial memory (Morris et al., 1982; Zola-Morgan and Squire, 1993), the computations that are performed by each of its subregions, particularly by the dentate gyrus (DG), are not well characterized. The sparse activity of the DG granule cells in both the number of active neurons and firing rate has motivated theories that DG facilitates memory formation through the distinct encoding of unique events. This is supported by the observation that spatial firing patterns of DG cells show pattern separation (Leutgeb et al., 2007; Neunuebel and Knierim, 2014; GoodSmith et al., 2019), which is in turn consistent with the general conceptual framework that the DG mossy fiber projections to CA3 support memory by promoting the generation of distinct hippocampal firing patterns for separate events (Marr and Brindley, 1971; Hopfield, 1982; Mcnaughton and Morris, 1987; Treves and Rolls, 1994). However, behavioral studies have suggested that the role of DG in memory may be broader, due to its essential role in support of complex spatial working memory (WM) where items need to be held in temporary storage for successful goal-directed behaviors (Xavier and Costa, 2009; Sasaki et al., 2018).

Consistent with a role of DG in working memory in addition to pattern separation, CA3 SWRs have also been shown to occur during WM, to depend on the DG during WM, and to predict subsequent correct choices (Sasaki et al., 2018). These findings suggest that the associative circuits created by the dense direct and indirect recurrent pathways in the DG-CA3 network (Amaral and Witter, 1989; Scharfman, 2007) support the generation of SWRs (150 – 250 Hz oscillation), which are thought to originate in CA3 with modulation from the CA2 and DG subregions (Buzsáki, 1986; Csicsvari et al., 2000; Oliva et al., 2016; Sasaki et al., 2018). SWRs occur during slow-wave sleep and during pauses in ongoing behavior and consist of short bouts of neuronal activity that correspond to a time-compressed replay of sequences from behavioral episodes (Lee and Wilson, 2002; Foster and Wilson, 2006; Pfeiffer and Foster, 2013; Joo and Frank, 2018). Decoding of such sequences in CA1 has revealed that they reflect available trajectories within an environment including spatial locations either behind or in front of the animal (Diba and Buzsáki, 2007; Gupta et al., 2010). Stored hippocampal sequences have in turn been shown to be important for learning and memory consolidation (Girardeau et al., 2009; Jadhav et al., 2012; Norimoto et al., 2018; Fernández-Ruiz et al., 2019) and are also proposed to guide behavior by facilitating decision making and future route planning (Gupta et al., 2010; Pfeiffer and Foster, 2013; Singer et al., 2013; Joo and Frank, 2018; but see Gillespie et al., 2021).

While the relation between SWRs and memory has been predominantly investigated in the hippocampal CA1 region, the finding that DG-dependent SWR activity in CA3 reflects the planning of future action (Sasaki et al., 2018) are consistent with a proposed broader role of DG in which its circuits are hypothesized to perform error correction to maintain accurate temporal order during the chaining of CA3 sequences during behavior (Lisman et al., 2005). However, complex memory tasks are highly dynamic with frequent transitions between brain states associated with predominant frequency ranges, each likely reflecting distinct underlying network mechanisms for memory encoding and retrieval (Kay and Frank, 2019). A complementary brain oscillation that is also strongly associated with sequential neuronal activity is theta (6-10 Hz oscillation). Unlike SWR activity that occurs during non-movement, theta is prominent and critical when memory is encoded during periods of active exploration (Buzsáki, 2005; Colgin, 2013; Quirk et al., 2021).

During theta states the binding of cells into sequences has been proposed to be supported by theta phase precession (O’Keefe and Recce, 1993; Dragoi and Buzsáki, 2006; Wang et al., 2015), which is a network mechanism that results in sequences when spiking of multiple overlapping place cells is organized within each theta cycle. For each place cell, spiking first occurs at a late theta phase upon entry into a place field and at progressively earlier phases of theta at the exit from the field. This mechanism organizes the sequential firing such that firing for place cells that are further ahead of the animal spike later within the theta cycle. The resulting compression of sequences from the behavioral timescale (seconds) to within the timescale of synaptic plasticity (milliseconds) may facilitate the storage of sequences in synaptic matrices (Bi and Poo, 1998; Melamed et al., 2004; Byrnes et al., 2011). At the population level, the precession of individual place cells can lead to spike sequences within each theta cycle (Dragoi and Buzsáki, 2006; Foster and Wilson, 2006; Feng et al., 2015; Chadwick et al., 2016) that may also reflect past and future paths of the animal (Johnson and Redish, 2007; Wang et al., 2020). Therefore, phase precession is a critical mechanism during theta states for organizing spike sequences during behavior and has been observed in the neural networks of all hippocampal subregions as well as in their theta modulated input, the medial entorhinal cortex (MEC) (Skaggs et al., 1996; Hafting et al., 2008).

While the DG has been found to be required for CA3 SWR generation and the retrieval of prospective representations in SWR sequences during slow wave activity (Sasaki et al., 2018), its role in supporting the precisely timed activity of CA3 neurons during theta states has not been explored. Here, we examined the contributions of DG inputs to CA3 theta phase precession to provide an understanding of the mechanisms of CA3 sequence generation across brain states. For comparison, we also examined the contribution of MEC inputs, which are the second major theta-modulated input to DG. To distinguish the role of the DG and MEC inputs to CA3 theta phase precession, we analyzed and compared CA3 network dynamics during theta from previously published recordings of CA3 cells during working memory tasks with either intact or diminished MEC or DG inputs (Sasaki et al., 2018; Sabariego et al., 2019). Based on our finding that DG contributes to prospective coding during SWRs (Sasaki et al., 2018), we hypothesized that DG also predominantly controls the emergence of prospective coding during theta oscillations. Our results are consistent with a contribution of DG, but not MEC inputs, to prospective coding and to the organization of temporal relations between CA3 neurons. We devised a phenomenological computational model to synthesize these findings and make predictions about inhibitory circuit elements that might support phase precession and sequence coding in CA3.

## Results

To test the contribution of DG inputs and of MEC inputs to CA3 phase precession, two previously published datasets with recordings of neuronal activity in the rat hippocampal CA3 region were analyzed (Sasaki et al., 2018; Sabariego et al., 2019). In these datasets, CA3 cells were recorded in hippocampus-dependent working memory tasks after lesioning either dentate granule neurons or the MEC (Fig. S1a,b). Each lesion group was paired with a respective control group (DG-lesioned and control: 16 sessions from 9 rats and 7 sessions from 4 rats; MEC-lesioned and control: 20 sessions from 8 rats and 18 sessions from 7 rats; Table S1). Because the working memory tasks require the rats to follow chosen trajectories, coverage of space was inevitably non-uniform. We therefore reasoned that the method of defining spike trains by first identifying place fields and then identifying passes through fields may be less precise as a result of uneven coverage. We therefore directly identified bouts of increased neuronal activity by selecting spike trains from the temporal firing patterns. We first selected candidate events based on the criterion that 5 or more spikes needed to occur with inter-spike intervals of less than 500 ms. We then determined behavioral measures, such as running speed and distance travelled from the corresponding time interval (Fig. S1d) and required a minimum distance (> 20 cm) and speed (> 2 cm/s) (see Fig. S1c for details on the criteria) to only consider neuronal activity during locomotion, when there is reliable occurrence of theta oscillations. Given that we included only spikes within trains and during movement periods in qualifying spike trains, only a proportion of all recorded spikes (34.3% vs. 20.4% DG control and lesion; 30.6% vs. 22.5% MEC control and lesion) were included in qualifying trains and further analyzed (Fig. S2a) for their timing with reference to the LFP signal. While spike trains were identified solely by timing and velocity criteria, we confirmed that spike trains clustered preferentially at one or few spatial locations, as would be expected for CA3 place cells (Fig. S2b-h).

### DG granule cell input was necessary for the expression of phase precession in CA3 neurons

Although it has long been known that there is substantial phase precession in DG cells (Skaggs et al., 1996), it is not clear whether DG inputs are necessary for phase precession in its direct target cells in CA3. We therefore compared phase precession in CA3 cells between DG-lesioned and control rats, which were trained to perform a dentate-dependent radial 8-arm maze WM task (Table S1) (Sasaki et al., 2018). We began by determining the level of phase precession in control CA3 cells and plotted the theta phase of all spikes of qualifying trains of a cell against the distance that the animal travelled from the beginning to the end of each train (“slope-by-cell” analysis). Phase precession was evident in the negative circular-linear regression slope between theta phase and the position along the trajectory where spikes occurred (Fig. 1a). We then performed the corresponding analyses in DG-lesioned rats. In rats with DG lesions, substantial loss of mossy fiber innervation was previously confirmed for all recording sites that are included in the analysis, and the extent of DG granule cell loss was previously quantified by scoring the remaining mossy fiber density at CA3 recording sites (Sasaki et al., 2018). Here, we combined CA3 cells from all recordings at sites with complete or partial mossy fiber loss (see Methods for a detailed description). In CA3 cells recorded at these sites, phase precession was less pronounced and more variable than in controls (Fig. 1b). The median circular-linear regression slope value in the slope-by-cell analysis was −149.4° for the control CA3 cells and −79.2° for CA3 cells from the DG-lesioned animals. Both medians were significantly negative (Fig. 1c; control: *n* = 84 cells, Sign Rank test; z-statistic = −6.19, signed rank = 396, *p* = 2.96 × 10^−10^; DG-lesioned: *n* = 68 cells, z-statistic = −2.63, signed rank = 742, *p* = 0.0043) though with a reduced slope of phase precession in the DG-lesion compared to the control group (Fig. 1c; z-statistic = − 3.63, rank sum = 5446, *p* = 2.84 × 10^−4^, Wilcoxon rank sum test). When considering the fraction of CA3 cells with negative compared to positive slopes, a lower proportion displayed negative slopes in lesioned compared to controls (88.1% in CTRL^(DG)^ vs. 69.1% in LESION^(DG)^; χ^2^ test for proportions, χ^2^ = 8.34, *p* = 0.0039). When adding the further condition that the slopes had to be not just negative, but also pass a significance criterion, the proportion of cells with significantly negative slopes also differed (Fig. 1c, shaded bars; 61.9% in CTRL^(DG)^ vs. 38.2% in LESION^(DG)^; χ^2^ test for proportions, χ^2^ = 8.43, *p* = 0.0037).

**Figure 1.**
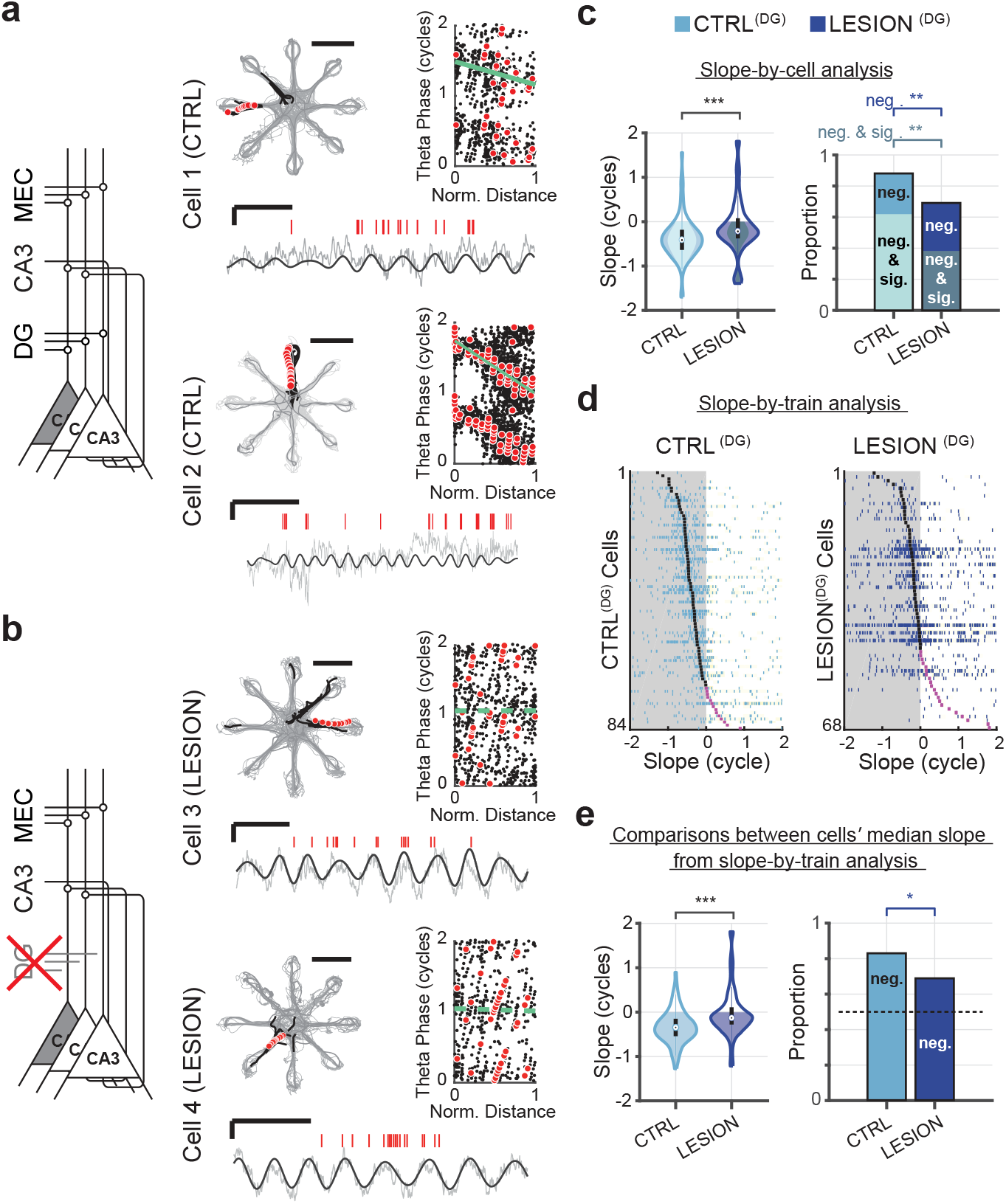
Dentate granule cell input is required for intact phase precession in CA3. **a**, Intact phase precession in control CA3 cells. Left, schematic of the major theta-modulated excitatory inputs to CA3. Right, Two example CA3 cells and their spatial firing patterns (grey lines, path; black dots, spike locations; red dots, spike locations of an example spike train; scale bar, 50 cm), an example spike train and corresponding LFP trace (red ticks, spikes; solid line, 6-10 Hz filtered LFP; gray line, raw LFP; scale bars, 250 ms and 500 μV), and phase-versus-normalized distance plot (black dots, spikes; red dots, spikes of example train, 1 cycle = 360°). Solid green lines in the plot indicate significant phase precession (*p* < 0.05). **b**, Data as in **a**, but for CA3 cells without DG inputs. Dashed green lines indicate the lack of phase precession. **c**, Phase precession slopes and proportions of phase precessing cells. One slope per cell was obtained by pooling the spikes of all trains and by fitting a circular-linear regression to this pool (slope-by-cell analysis). The median magnitude of the slopes (violin plots; *n* = 84 control (CTRL) and 68 DG lesion (LESION) CA3 cells, z-statistic = −3.62, rank sum = 5446, *p* = 2.8 × 10^−4^, Wilcoxon rank sum test) and the proportion of negative slopes (bar plots; only negative slopes, χ^2^ = 8.34, *p* = 0.0039; negative and significant slopes, χ^2^= 8.43, *p* = 0.0037, chi-square test) were reduced by the DG lesion. **d**, Slopes for all trains from control (CTRL^(DG)^, left) and DG-lesioned (LESION^(DG)^, right) CA3 cells (slope-by-train analysis). Each row depicts the slope values from each of the trains of one cell (blue ticks), and cells are sorted from top to bottom by their mean slope (black tick when negative, purple tick when positive). Shaded regions correspond to negative values. **e**, Phase precession slopes (violin plots; *n* = 84 control and 68 DG lesion CA3 cells, z-statistic = − 4.39, rank sum = 5242, *p* = 1.2 × 10^−5^, Wilcoxon rank sum test) and proportions of phase precessing cells (bar plots; χ^2^= 4.28, *p* = 0.038, chi-square test) from the slope-by-train analysis. For analysis of proportions, a cell was considered phase precessing if the median slope was negative. Violin plots in panels c and e: Outline, distribution; shading, negative slopes (inner shading in c, negative and significant slopes); error bars, 1.5 times the interquartile interval above the third and below the first quartile. * *p* < 0.05, ** *p* < 0.01, *** *p* < 0.001.

Differences between the control and DG-lesion groups were also apparent from the distribution of slopes obtained from the circular-linear regression analysis of single pass data (“slope-by-train” analysis; Fig. 1d). Here, we used the slopes of individual spike trains and, for statistical comparisons, averaged the slopes of each cell’s trains (Fig. 1e). We then compared the cells’ averages across groups and found that the median of cell-averaged slopes was significantly less than zero in control and DG lesioned rats, although the difference was less pronounced in the lesion group (CTRL^(DG)^: *n* = 84 cells, z-statistics = −6.3, signed rank = 381, *p* = 1.9 × 10^−10^, LESION^(DG)^: *n* = 68 cells, z-statistics = −1.98, signed rank = 849, *p* = 0.024). In addition, the median of cells from lesioned rats was significantly different from the median of control cells (z-statistic = −4.39, rank sum = 5242, *p* = 1.2 × 10^−5^, Wilcoxon rank sum test). Further, the proportion of CA3 cells with negative mean slopes was higher in cells from control than from lesioned rats (Fig. 1e, right; 83.3 % vs. 69.1 %, χ^2^ = 4.3, *p* = 0.038, chi-square test of proportions). Taken together, these analyses demonstrate that CA3 phase precession is diminished when the dentate granule cell input to CA3 neurons is reduced. In particular, the analyses with single train slopes revealed that the remaining inputs to CA3 after DG lesions are not sufficient to sustain the reliable, single-train phase precession thought to be required for real-time encoding and retrieval of episodic memories.

### MEC inputs to CA3 were also necessary for the expression of phase precession in CA3 neurons

The MEC is known to be necessary for CA1 phase precession (Schlesiger et al., 2015). However, it is not known whether CA3 also requires MEC input to generate phase precession or can generate phase precession with DG connectivity alone. Thus, we next tested whether DG alone can support CA3 phase precession by analyzing recordings of CA3 cells in MEC-lesioned rats. The MEC lesions were consistent between rats and included 93.0% of the total volume, with damage approximately matched across cell layers (95.3% of layer II, 92.4% of layer III, and 91.4% of deep layers) (Sabariego et al., 2019). We extracted qualifying spike trains recorded in CA3 of MEC-lesioned animals as described above (Table S1). The slope-by-cell analysis revealed that the CA3 cells of control rats displayed phase precession (Fig. 2a) and that precession was reduced but not abolished in MEC-lesioned animals (Fig. 2b and c; CTRL^(MEC)^: *n* = 101 cells, median slope: −120.4°; LESION^(MEC)^: *n* = 158 cells, −66.9°; control vs. lesioned: z-statistic = −2.34, rank sum = 11754, *p* = 0.0193, Wilcoxon rank sum test; control less than zero: z-statistic = −6.18, signed rank = 750, *p* = 3.16 × 10^−10^; lesioned less than zero: z-statistic = −3.6, signed rank = 4204, *p* = 1.57 × 10^−4^; sign tests). The proportion of cells with negative slopes was lower in the MEC-lesioned rats when all cells with negative slopes (83.2% and 70.3%, CTRL^(MEC)^ vs. LESION^(MEC)^; χ^2^ = 5.52, *p* = 0.0188) and when only cells with significantly negative slopes were considered (54.5% in CTRL^(MEC)^ vs. 38.6% in LESION^(MEC)^, χ^2^ = 6.26, *p* = 0.0124; χ^2^ tests for proportions). As with DG lesions, the slope-by-train analysis showed that reliable negative single train slopes were seen for trains from control CA3 cells but not for trains from CA3 cells of MEC-lesioned rats (Fig. 2d, e; Medians less than zero: CTRL^(MEC)^, z-statistic = −6.9, signed rank = 550, *p* = 3.4 × 10^−12^; LESION^(MEC)^, z-statistic = −2.9, signed rank = 4628, *p* = 0.0021; Median of CTRL^(MEC)^ vs. LESION^(MEC)^: z-statistic = −3.9, rank sum = 10832, *p* = 9.3 × 10^−5^, Wilcoxon rank sum test; proportions of negative slopes: 84.2% and 65.8%, CTRL^(MEC)^ vs. LESION^(MEC)^, χ^2^ = 10.5, *p* = 0.0012, chi-square test). These observations support a role for MEC in the generation of robust phase precession in the CA3 of rats. Therefore, the DG-CA3 network alone is incapable of generating phase precession at control levels—for this, both the DG and MEC inputs are necessary.

**Figure 2.**
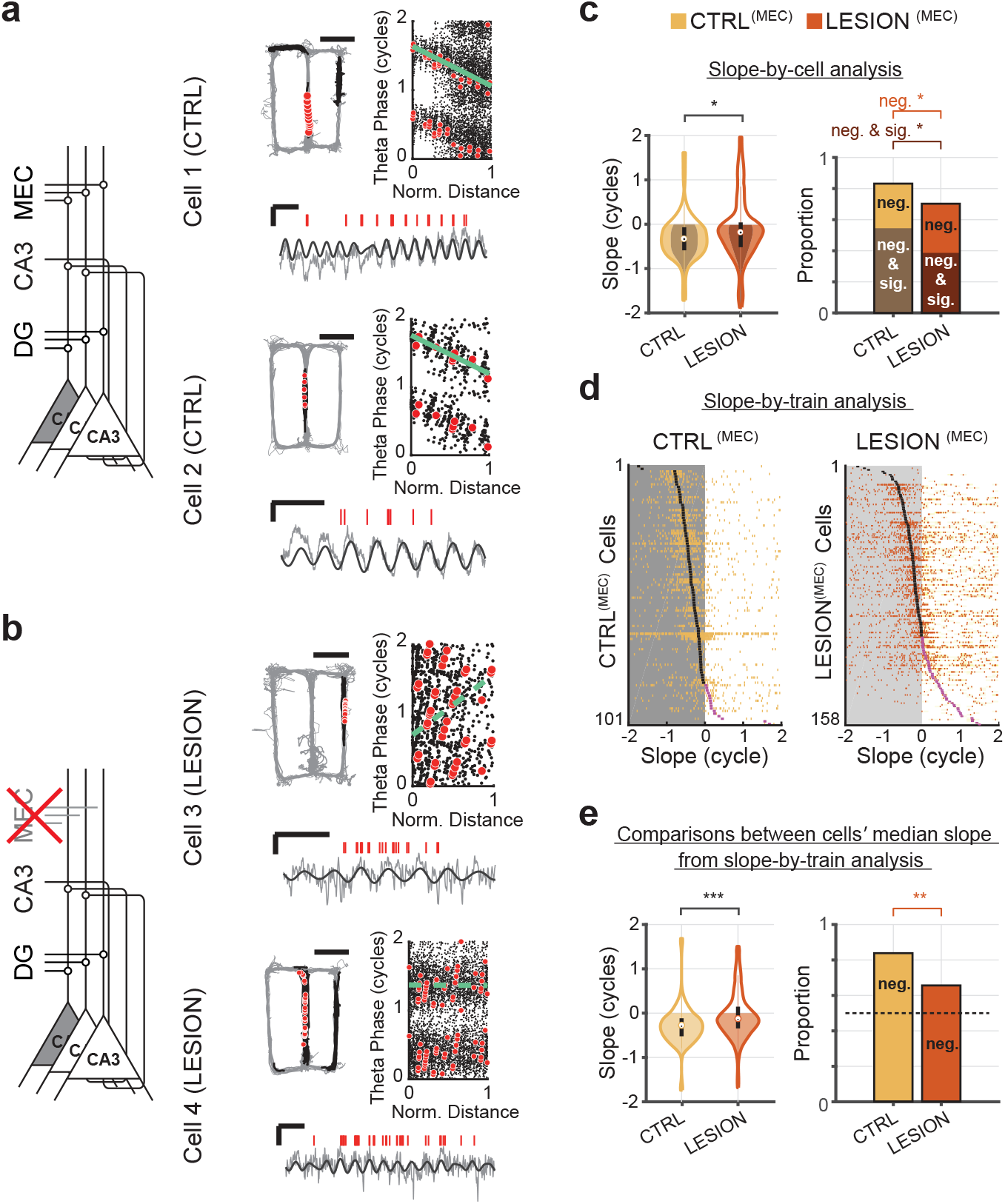
Medial entorhinal cortical input is required for intact phase precession in CA3. This figure follows the presentation of Fig. 1, but with data from CA3 cells with lesioned MEC inputs and respective controls. **a**, **b**, Firing patterns of example CA3 cells in control and MEC-lesioned rats. Solid green lines in the phase-versus-normalized distance plot indicate significant phase precession (*p* < 0.05) and stippled green lines indicate a lack of significant phase precession. Scale bars for LFP: 500 μV and 250 ms, for path: 50 cm. **c**, Phase precession slopes (violin plots; *n* = 101 control (CTRL) and 158 MEC lesion (LESION) CA3 cells, z-statistic = - 2.34, rank sum = 11754, *p* = 0.019, Wilcoxon rank sum test) and proportions of phase precessing cells (bar plots; controls vs. lesion, only negative slopes, χ^2^= 5.52, *p* = 0.019; negative and significant, χ^2^= 6.26, *p* = 0.0124, chi-square test) from the slope-by-cell analysis. The magnitude of the slopes and the proportion of negative slopes were reduced by the MEC lesions. **d**, Slopes for all trains from CTRL^(MEC)^ (left) and LESION^(MEC)^ (right) CA3 cells. Each row depicts the slope values from each of the trains of one cell (yellow and orange ticks), and cells are sorted from top to bottom by their median slope (black tick when negative, purple tick when positive). **e**, Phase precession slopes (violin plots; *n* = 101 control and 158 MEC lesion CA3 cells, z-statistic = −3.91, rank sum = 10832, *p* = 9.3 × 10^−5^, Wilcoxon rank sum test) and proportions of phase precessing cells (bar plots; χ^2^ = 10.50, *p* = 1.2 × 10^−3^; chi-square test) from the slope-by-train analysis. Violin plots in panel c and e: Outline, distribution; shading, negative slopes (inner shading in c, negative and significant slopes); error bars, 1.5 times the interquartile interval above the third and below the first quartile. * *p* < 0.05, ** *p* < 0.01, *** *p* < 0.001.

### DG and MEC lesions had qualitatively distinct effects on CA3 phase precession

After confirming that both the DG and MEC were necessary for CA3 phase precession at control levels, we asked whether there were qualitative differences in the phase precession patterns when each of these inputs were diminished. Phase precession can be reduced by either limiting the theta phase range over which spiking occurs or by heightening the variability around a monotonically decreasing precession slope, or both. To determine whether the theta phase range was altered by the lesions, we calculated the onset and offset theta phase of CA3 spike trains. The onset phase of trains – defined as the circular median phase of the spikes in the first cycle of each train – showed a marked shift toward earlier phases in CA3 cells of DG-lesioned rats compared to control rats (Fig. 3a). For control cells, the peak of the distribution of onset phases (Φ_on_) of all trains was during the late phase of the theta cycle, past the trough, and accordingly, the circular mean of all onset phases was 245.4°. For cells from DG-lesioned rats, the circular mean of all onset phases was shifted towards the trough (223.3°, *n* = 1,548 and 1,354 spike trains for control and lesion, respectively; test statistic = 420.3, *p* = 5.4 × 10^−92^, circular MANOVA, but even more strikingly, was more broadly and symmetrically distributed around the trough (onset phase concentration parameters: CTRL^(DG)^ κ = 1.48, LESION^(DG)^ κ = 0.26, U-statistic = 415.7, *p* = 2.1 × 10^−92^, concentration test). As expected for phase precessing cells, the distributions of offset phases (Φ_off_) – defined as the circular median phase of the spikes in the last cycle of each train – was earlier in the theta cycle for cells from control and DG-lesioned rats with mean offset phases of 81.7° and 60.1°, respectively. The offset phase thus shifted to a similar degree as the onset phase in DG-lesioned rats compared to control rats (Fig. 3b; test statistic = 21.1, *p* = 2.6 × 10^−5^, circular MANOVA), but the change in dispersion between controls and DG-lesioned rats was more modest than for the onset phase (offset phase concentration parameters: CTRL^(DG)^ κ = 0.41, LESION^(DG)^ κ =, 0.59, U-statistic = 11.3, *p* = 7.8 × 10^−4^, concentration test). Furthermore, effects on CA3 spike timing were more pronounced with DG lesions compared to MEC lesions (Fig. 3c,d). The onset phase of the MEC-lesioned CA3 trains showed only a minor difference from controls (Fig. 3c; CTRL^(DG)^ 240.5°, LESION^(DG)^ 246.9°, *n* = 2,326 and 3,132 spike trains for control and lesion, respectively; test statistic = 29.6, *p* = 3.7 × 10^−7^, circular MANOVA) and only slight broadening in MEC-lesioned compared to control rats (CTRL^(MEC)^ κ = 0.95, LESION^(MEC)^ κ = 0.74, U-statistic = 22.9, *p* = 1.7 × 10^−6^, concentration test). For the offset phase, neither the mean nor the concentration differed between MEC lesion and control cells (Fig. 3d; mean, CTRL^(MEC)^: 76.6° LESION^(MEC)^ 84.8°, test statistic = 4.1, *p* = 0.129, circular MANOVA, concentration: CTRL^(MEC)^ κ = 0.53, LESION^(MEC)^ #x03BA; =, 0.55, U-statistic = 0.12, *p* = 0.73, concentration test). These results indicate that it is predominantly the DG input rather than the MEC input to CA3 that is involved in setting the theta phase of CA3 spikes, in particular the precise onset phase within the late phase of the theta cycle. This effect is consistent with the hypothesis that DG inputs are essential for generating intrinsic “look-ahead” spikes in CA3 (Hasselmo et al., 2002; Lisman et al., 2005; Sanders et al., 2015; Yiu et al., 2022).

**Figure 3.**
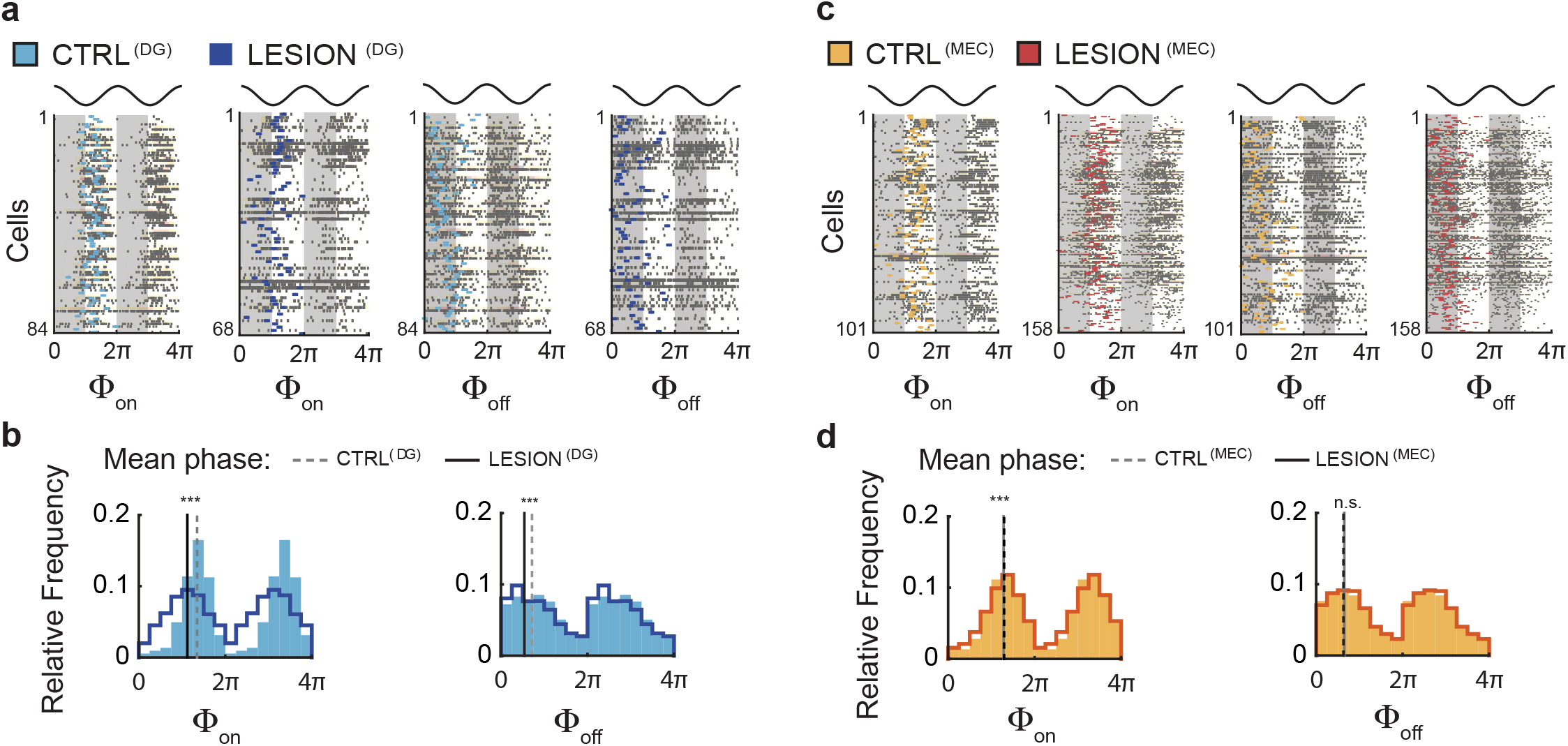
DG lesions, but not MEC lesions substantially broaden the onset phase of CA3 spike trains. **a**, Onset and offset phases of spike trains from CA3 cells in control (CTRL^(DG)^) and DG-lesioned rats (LESION^(DG)^). In each of the four raster plots, each row displays the onset phase (Φ_on_) or offset phase (Φ_off_) of the train of a cell (in grey), and the cell’s median onset phase (light blue, control; dark blue, DG-lesioned). Data are repeated from 2π to 4π for clarity, and two theta cycles are displayed on top for reference. **b**, Onset and offset phases are compared between spike trains from control and DG-lesioned rats. Data are repeated from 2π to 4π for clarity. Note the clustering of onset phases in the ascending portion of the theta cycle in control spike trains in contrast to the greater dispersion of onset phases spike trains from DG-lesioned rats. Onset phases shifted by an average of −22.1° between CTRL^(DG)^ (light blue bars) and LESION^(DG)^ groups (dark blue line; *n* = 1,548 and 1,354 spike trains, test statistic = 420.3, *p* = 5.4 × 10^−92^, circular MANOVA). In addition, the lesion resulted in a greater dispersion of onset phases over the entire theta cycle (onset phase concentration: CTRL^(DG)^ κ = 1.48, LESION^(DG)^ κ = 0.26, U-statistic = 415.7, *p* = 2.1 × 10^−92^, concentration test). Offset phases were skewed towards the descending portion of the theta cycle and were shifted by an average of −21.6° by the lesion (test statistic = 21.1, *p* = 2.6 × 10^−5^, circular MANOVA). Compared to the pronounced broadening of onset phases by the lesion, the effect on the dispersion of the offset phases was more minor, albeit statistically different (offset phase concentration: CTRL^(DG)^ κ = 0.41, LESION^(DG)^ κ = 0.59, U-statistic = 11.3, *p* = 7.8 × 10^−4^, concentration test). **c-d**, As (a-b), but for the cells from MEC-lesioned rats (LESION^(MEC)^) and their corresponding controls (CTRL^(MEC)^). Onset phase values shifted a modest −6.4° along with minor sharpening of the dispersion by the lesion (*n* = 2,326 CTRL^(MEC)^ and 3,132 LESION^(MEC)^ spike trains, phase: test statistic = 29.6, *p* = 3.7 × 10^−7^, circular MANOVA; concentration: CTRL^(MEC)^ κ = 0.95, LESION^(MEC)^ κ = 0.74, U-statistic = 22.9, *p* = 1.7 × 10^−6^, concentration test). The mean and the dispersion of offset phases were not different between cells of MEC-lesioned and control rats (phase difference = −8.2°, test statistic = 4.1, *p* = 0.129, circular MANOVA; concentration: CTRL^(MEC)^ κ = 0.53, LESION^(MEC)^ κ = 0.55, U-statistic = 0.12, *p* = 0.73, concentration test). n.s., not significant, *** *p* < 0.001.

The finding that the phase shift of the onset phase is coupled with a broadening of the phase dispersion can be interpreted as the emergence of “noise” spikes in early theta phases which would increase the phase variance and shift the mean to earlier phases upon entry to the place field. To further examine this possibility, we analyzed several measures of spike phase distribution. When measuring spike phase distribution across the theta cycle regardless of the distance travelled by the rat, CA3 spike phase in control cells was concentrated in the middle of the theta cycle, as expected (Fig. 4a; Buzsaki et al., 1983; Fox et al., 1986). Spike phase distributions across the theta cycle shifted towards earlier phases of the theta cycle with DG lesions while the shift was in the opposite direction with MEC lesions compared to their respective controls (Fig. 4a; mean theta phase: CTRL^(DG)^ *n* = 27,988 spikes, mean phase = 183.2°, LESION^(DG)^ *n* = 28,704 spikes, mean phase = 70.5°, χ^2^ statistic = 973.199, *p* = 4.7 × 10^−212^; CTRL^(MEC)^ *n* = 62,294 spikes, mean phase = 147.7°, LESION^(MEC)^ *n* = 68,385 spikes, mean phase = 164.9°, χ^2^ statistic = 191.408, *p* = 2.7 × 10^−42^; χ^2^ test for proportion of spikes contained in each of the three theta bins). The shift to earlier phases was accompanied by an increase in the proportion of spikes in the first third of the theta cycle in DG-lesioned animals, which could result either from the addition of spurious spikes compared to controls or solely from redistributing the same number of spikes over a broader phase. To test whether additional spikes were present in early theta cycles in DG lesioned rats, we counted the number of spikes in the first full theta cycle, and for each neuron, averaged the number of spikes over all spike trains (Fig. 4b). In the DG-lesioned rats, a median number of 1.28 spikes occurred in the first full theta cycle compared to 0.54 spikes in control rats (z-statistic = −3.63, rank sum = 5446, *p* = 2.8 × 10^−4^; Wilcoxon rank sum test). In MEC-lesioned rats, a median of 1.42 spikes occurred in this cycle compared to a median 1.18 spikes in control rats (z-statistic = −2.03, rank sum = 11938.5, *p* = 0.043; Wilcoxon rank sum test). Thus, the DG lesions resulted in a ∼2.4-fold increase in the spike counts in the initial theta cycle compared to a ∼1.2-fold increase induced by MEC lesions, supporting the idea that spurious spikes are added to early cycles in addition to a general broadening of the onset phase distribution (Fig. 3b).

**Figure 4.**
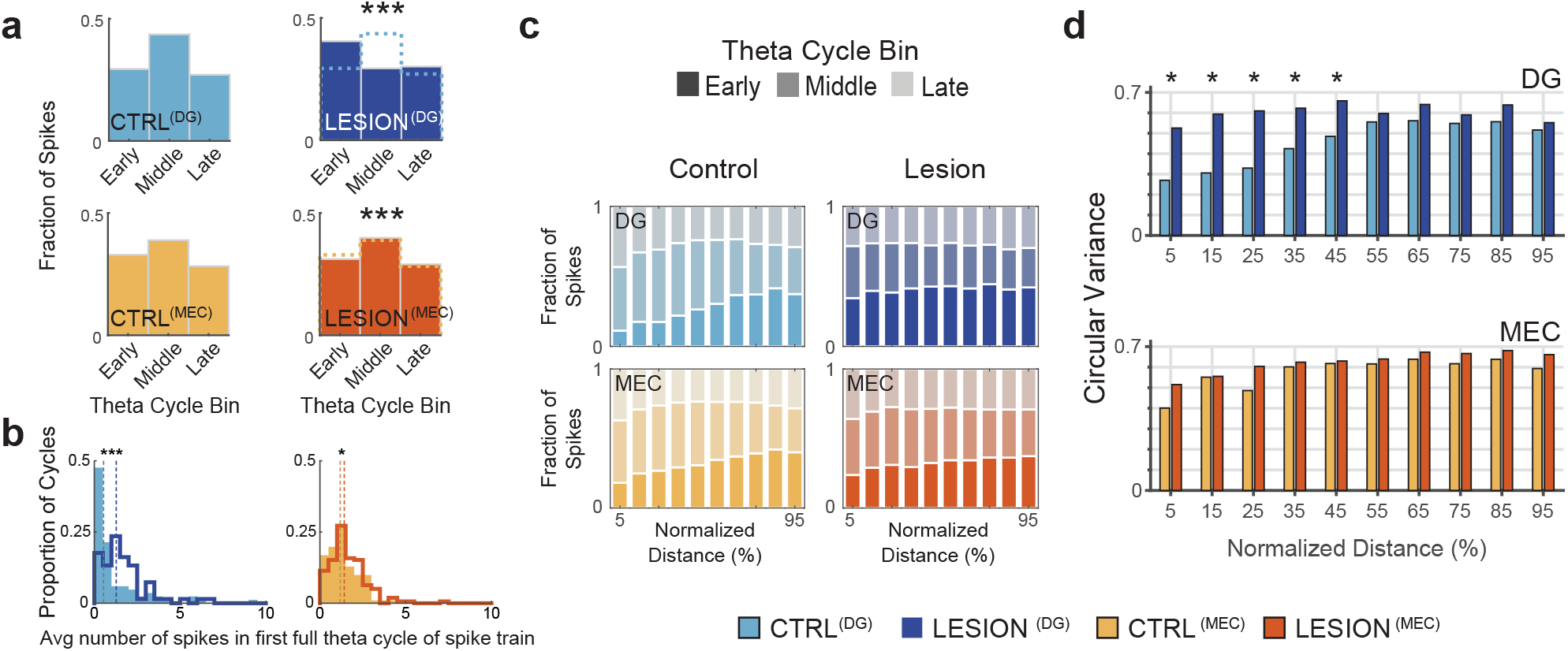
With reduced DG input, peak probability of CA3 spiking shifts to earlier theta phases. **a**, Left, For control CA3 cells, distribution of spike incidence across early, middle, and late theta phases. The middle of the theta cycle contains more spikes than early or late phases. Top right, The proportion of CA3 spikes early in the theta cycle was markedly increased with reduced DG granule cell input (LESION^(DG)^; *n* = 27,988 CONTROL^(DG)^ spikes and 28,704 LESION^(DG)^ spikes, χ*^2^*= 973.2, *p* = 4.7 × 10^−212^, χ^2^ test with control vs. lesion and theta phase as factors). Dotted lines, data from control CA3 cells, as shown to the left. Bottom right, with reduced MEC inputs (LESION^(MEC)^), there was a small shift for CA3 spikes to occur later in the theta cycle (*n* = 62,294 CONTROL^(DG)^ spikes, 68,385 LESION^(DG)^ spikes, χ*^2^*= 191.4, *p* = 2.7 × 10^−42^, χ^2^ test). **b**, Average number of spikes in the first full theta cycle of spike trains. In trains from DG-lesioned rats, the median number of spikes increased ∼2.4 fold compared to control trains (from 0.54 to 1.28, *n* = 84 CONTROL^(DG)^ cells, *n* = 68 LESION^(DG)^ cells, z-statistic = - 3.63, rank sum = 5446, *p* = 2.8 × 10^−4^; Wilcoxon rank sum test). In MEC-lesioned rats, the median number of spikes increased 1.2-fold compared to control trains (from 1.18 to 1.42 spikes, *n* = 101 CONTROL^(MEC)^ cells, *n* = 158 LESION^(MEC)^ cells, z-statistic = −2.03, rank sum = 11938.5, *p* = 0.043; Wilcoxon rank sum test). **c**, Fraction of CA3 spikes at early, middle, and late phases of the theta cycle as a function of normalized (%) distance during the train. At the onset of control trains, a low proportion of spikes is typically observed at early phases, but this proportion increased with DG lesions. **d**, Top, Consistent with the addition of early-phase spikes upon entry into the place field, DG lesions selectively increased the variability of spike phase in the first half of a field traversal (from 5% to 45% of normalized distance, Wilcoxon rank sum tests; Holm-Bonferroni corrected; see Table S2 for statistics). Bottom, MEC lesions did not increase the variability of theta phase along the distance through the field. (Wilcoxon rank sum tests; Holm-Bonferroni corrected; Table S2). * *p* < 0.05, *** *p* < 0.001.

In additional analyses that consider the distance travelled during a spike train, we binned the normalized distance during each spike train into 10 bins and considered the joint distribution of spike phase and normalized distance (Fig. 4c). Here, we found that DG lesions resulted in a particularly pronounced redistribution of spikes to earlier theta phases in early bins. This redistribution reduced the proportion of late phase spikes that are normally observed in the early section of the path such that the phase when spikes occurred was now remarkably similar irrespective of distance along the path. In addition, the circular variance of CA3 spike theta phase was selectively increased in the first half of the path (Fig. 4d, top; pairwise comparisons between CTRL^(DG)^ and LESION^(DG)^ are significant in the first five of ten bins, Holm-Bonferroni corrected *p* < 0.05, see Table S2 for detailed statistics) such that the variance in all bins with lesion reached levels that are in controls only observed in the second half. In contrast, circular variance took on similar values in MEC control and lesioned rats (Fig. 4d, bottom; pairwise comparisons between CTRL^(MEC)^ and LESION^(MEC)^ all bins not significant, see Table S2 for detailed statistics). The broadening of the onset phase distribution at the onset of the train together with the increase in firing rate in the initial theta cycle (Fig. 4c) are consistent with the possibility that DG to CA3 input is critically involved in restricting the spiking in the beginning of the train to late-theta phase (“prospective”) CA3 spiking by inhibiting spurious spikes in the early phases of the theta cycle.

### Putative granule cells exhibited a narrow theta phase preference at the onset of spiking

To next ask whether the temporal profile of DG granule cell spiking is precise enough to organize the spiking phase of CA3 neurons, we analyzed neuronal activity from rats in which we were able to record single units from the DG (*n* = 5 cells, see Methods). Putative granule cells showed phase precession (Fig. S3a,b), which was accompanied by a strikingly narrow theta phase preference at the onset of spiking. The phase preference then broadened over the course of the spike train (Fig. S3c). Putative mossy cells and CA3 pyramidal cells, although also phase precessing, showed a relatively broad theta phase variability throughout the entire spike train (Fig. S3c). These findings are complementary to the results from the DG lesion experiments, and consistent with a particularly critical role of DG granule cells for providing temporal information to CA3 pyramidal cells upon entering the place field.

### Analytically reducing the phase variability partially recovered phase precession of CA3 cells from MEC-lesioned rats

If a lesion impairs phase precession by increasing the variance of the phase distribution, it might be feasible to recover phase precession by analytically reducing the phase variance. Phase variance can be reduced by replacing, within each theta cycle, all spikes with their mean timestamp. However, if the main effect of a lesion is a reduction in slope or phase range, replacing the spikes with their cycle mean should not restore phase precession. When we replaced the spikes of each theta cycle with their mean phase within the cycle (‘cycle-mean analysis’), the effect of the lesion could be largely recovered in MEC-lesioned rats, but not in DG-lesioned rats. The rescue of the MEC lesion was observed both when all trains were combined as well as when individual trains were analyzed separately (Fig. S4). With the cycle-mean analysis, the mean of the cells’ slopes in MEC lesioned rats was no longer distinguishable from control rats (CTRL^(MEC)^: −97.4° per traversal vs. LESION^(MEC)^: −84.6° per traversal, z-statistic = −1.44, rank sum = 12285, *p* = 0.151, Wilcoxon rank sum test), while it remained different from controls in DG-lesioned rats (CTRL^(DG)^: −96.6° per traversal vs. LESION^(DG)^: −5.6° per traversal, z-statistic = −4.23, rank sum = 5285, *p* = 2.4 × 10^−5^, Wilcoxon rank sum test) (Fig. 5a-c). In all cases, however, the mean slope was less than zero for each of the groups (CTRL^(DG)^: *n* = 84 cells, z-statistic = −6.49, *p* = 4.4 × 10^−11^, LESION^(DG)^: *n* = 68 cells, z-statistic = −2.33, *p* = 0.01; CTRL^(MEC)^: *n* = 101 cells, z-statistic = −7.42, *p* = 5.9 × 10^−14^, LESION^(MEC)^: *n* = 158 cells, z-statistic = −7.56, *p* = 2 × 10^−14^; sign tests), as without cycle averaging (Fig. 5c; compare with Figs. 1 and 2). We also tested to what extent the calculation of the cycle mean reduced phase variability compared to the phase variability in the original spike trains. In DG-lesioned rats, there was only a minor difference in the variance of the phase probability over all trains of each neuron between the original and cycle-mean analyses (circular variance = 0.733, cycle-mean circular variance = 0.661, z-statistic = 2.04, *p* = 0.041, Wilcoxon rank sum test; Cohen’s *d* = − 0.308). However, in the two control groups and in the MEC-lesion group variance was substantially reduced by taking the cycle mean (CTRL^(DG)^: circular variance = 0.707, cycle-mean circular variance = 0.557; CTRL^(MEC)^: circular variance = 0.782, cycle-mean circular variance = 0.603; LESION^(MEC)^: circular variance = 0.805, cycle-mean circular variance = 0.634; tests of difference between original and cycle-mean variances: all *p* values < 10^−6^, Wilcoxon rank sum tests; Fig. 5d; compare, for each panel, the difference of medians indicated by the solid and dashed vertical lines; Cohen’s *d* for CTRL^(DG)^, CTRL^(MEC)^, and LESION^(MEC)^, respectively: - 0.809, −0.917, −0.779). Taken together, these results indicate that increased within cycle variability makes a major contribution to diminished phase precession in cells from MEC-lesioned animals. In contrast, increased within cycle variability was not identified as the key source for reducing phase precession in DG lesioned animals. The mechanisms by which DG and MEC circuits contribute to the organization of the precise temporal profiles of CA3 spiking (Fig. 5e-f) are therefore distinct and are consistent with a model in which DG inputs effectively restrict spiking to late phases of the theta cycles early in the spike train of a CA3 neuron (i.e., upon entry into the place field) (Fig. 5e). In contrast, later in the spike train (i.e., in the middle and near the exit from the place field), MEC inputs appear to ensure an appropriate mean theta phase of CA3 spiking by driving CA3 neurons in time windows around a monotonically decreasing mean theta phase over successive theta cycles (Fig. 5f).

**Figure 5.**
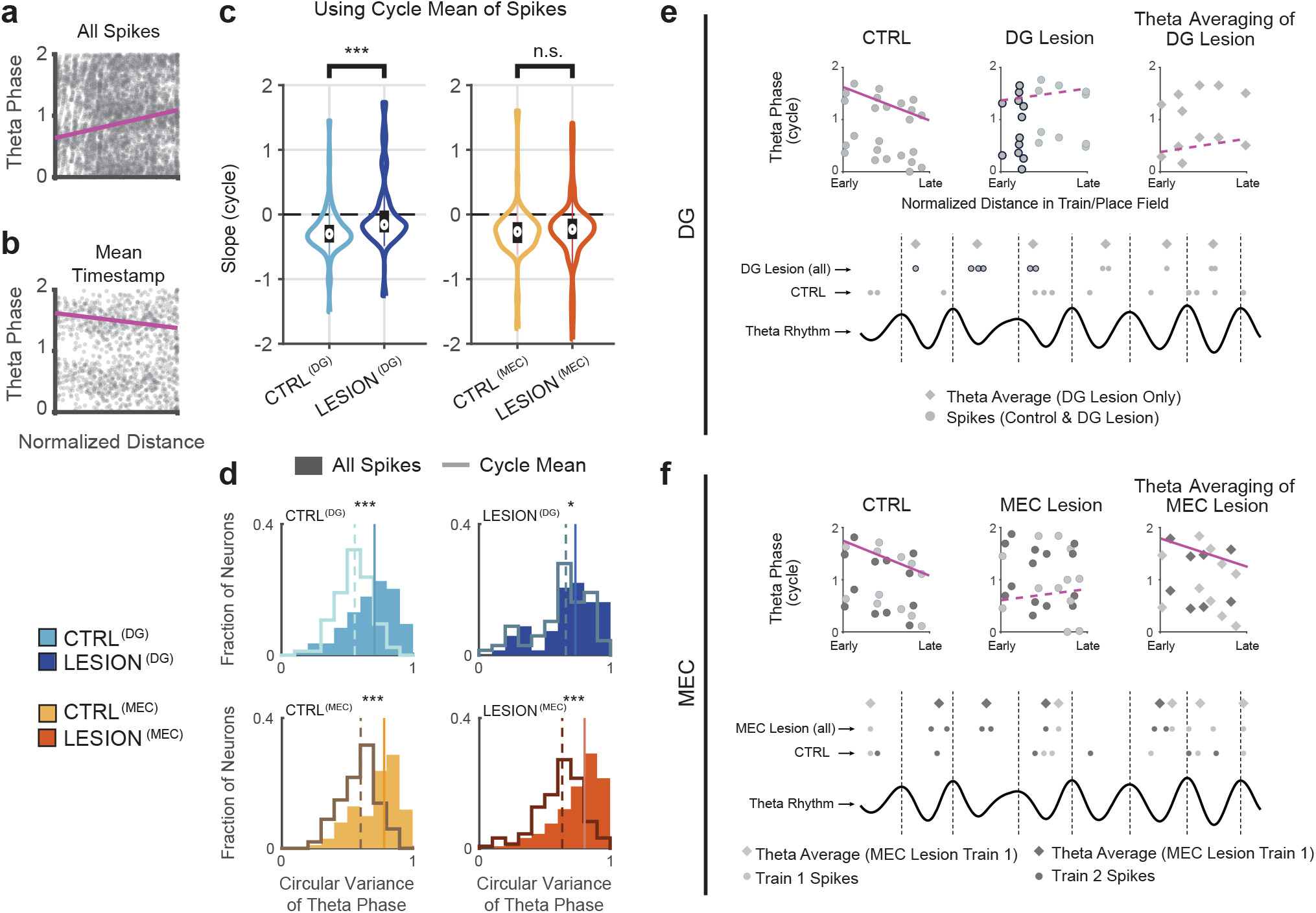
Distinct patterns of temporal reorganization of CA3 spiking after DG and MEC lesions. **a**, Phase-distance plots of spikes from an example CA3 cell from an MEC-lesioned rat. All spikes of the cell’s trains are included. **b**, Phase-distance plot of the same cell after replacing the spikes within each theta cycle with the mean phase of the spikes within each cycle. Using the cycle mean yields a negative precession slope. **c**, Distribution (violin plot) of circular-linear regression slopes calculated from the cycle means. Using the regression slopes from cycle means did not result in a difference from controls for CA3 cells of MEC-lesioned rats (z-statistic = - 1.44, *p* = 0.151, Wilcoxon rank sum test), while the difference was retained for cells of the DG-lesioned rats (z-statistic = −4.23, *p* = 2.4 × 10^−5^, Wilcoxon rank sum test). However, as expected from previous analyses without phase averaging, the median slope was less than zero for each of the groups (CTRL^(DG)^: *n* = 84 cells, z-statistic = −6.49, *p* = 4.4 × 10^−11^, LESION^(DG)^: *n* = 68 cells, z-statistic = −2.33, *p* = 0.01; CTRL^(MEC)^: *n* = 101 cells, z-statistic = −7.42, *p* = 5.9 × 10^−14^, LESION^(MEC)^: *n* = 158 cells, z-statistic = −7.56, *p* = 2 × 10^−14^; sign tests), as without cycle averaging (see Figs. 1 and 2). A broader distribution of spikes within theta-cycles after MEC lesions may have precluded the detection of phase precession. **d**, Circular variance of theta phase with either all spikes (filled bars) or with each cycle’s spike mean (solid lines). In all groups, the circular variance decreased after replacing each cycle’s spikes with their mean, though the effect is least pronounced for CA3 cells of DG-lesioned rats. **e**, Schematic of how DG lesions can disrupt CA3 temporal coding. Top left, In control animals, initial spikes of the place field occur late in the theta cycle and progressively advance to earlier phases further into the place field. Spikes are depicted by the gray circles. Top middle, Compared to the control condition, DG lesions result in additional CA3 spikes early in the theta cycle at the onset of trains (gray circles with black outline). Later spikes in the middle and end of the place field remain largely intact. Top right, theta cycle averaging (gray diamonds) is unable to rescue phase precession because the slope remains reduced after this calculation. Bottom, Schematic of spiking with reference to the theta cycle, corresponding to the above plots. **f**, Schematic of how MEC lesions can disrupt CA3 temporal coding. Top left to right, Two spike trains of cells are depicted by different shades of gray. Increased spike time variability within and across spike trains in each cycle lowers the likelihood of detecting phase precession with MEC lesions in the spike-by-cell analysis. Theta averaging rescues phase precession because, in contrast to (e), it at least partially reverses the heightened variability across successive traversals of the place field (i.e., trains 1 and 2). MEC thus supports the consistency of theta phase coding in CA3 pyramidal cells. Violin plots: Outline, distribution; error bars: 1.5 times the interquartile interval above the third and below the first quartile. n.s., not significant, * *p* < 0.05, *** *p* < 0.001.

### Theta-scale temporal correlations of CA3 cells were preserved with MEC lesions but not with DG lesions

Phase precession is associated with the occurrence of ordered neuronal firing patterns within each theta cycle (Dragoi and Buzsáki, 2006) such that sequential firing within a theta cycle (‘theta sequence’) corresponds to the order in which place fields are traversed, but with the timing in the theta cycle compressed compared to the behavioral time scale. For example, two adjacent place cells are activated one after another within milliseconds in a theta cycle, while the rise and fall in firing rates when traversing the fields occurs on a much slower time scale. Phase precession is thought to link the slower behavioral sequence to the faster pairwise temporal correlation in theta cycles. Because phase precession in a novel environment can be observed earlier than theta sequences, it has been proposed that intact phase precession is a prerequisite for the emergence of precise timing on the theta scale (Feng et al., 2015). However, it is not known whether there is indeed a definite relation such that theta sequences can only emerge in conditions with intact phase precession (Foster and Wilson, 2007). Here we therefore asked whether the precise pairwise timing at the theta scale was diminished when phase precession was impaired by the reduction of either DG or MEC inputs to CA3. To measure whether time-compression occurred, we measured whether there is a relation between the timing on the theta scale (i.e., the temporal difference in the spike cross-correlation of cell pairs) and the behavioral scale (i.e., the distance between the peak firing locations of place fields; see Methods) (Dragoi and Buzsáki, 2006).

While there was a strong correlation between the spatial separation of place fields and the theta phase difference of cell pairs in controls (CTRL^(DG)^: *n* = 30 pairs, r = 0.651, *p* = 9.87 × 10^−5^, Pearson’s correlation), we found that the behavioral order of firing was not reflected on the theta-cycle time scale in cell pairs from DG-lesioned rats (Fig. 6a-d; LESION^(DG)^: *n* = 14 pairs, r = - 0.187, *p* = 0.523, Pearson’s correlation). This effect was observed even though the proportion of neuron pairs with phase precession in DG-lesioned rats was comparable to that in control rats (Fig. 6d, right). Contrary to the result with DG lesions, the CA3 pairs in the MEC-lesioned rats maintained their spiking order in theta cycles compared to their firing order on the maze (Fig. 6e-h; CTRL^(MEC)^ *n* = 19 pairs, r = 0.654, *p* = 0.0024; LESION^(MEC)^ *n* = 27 pairs, r = 0.624, *p* = 5.02 × 10^−4^; Pearson’s correlations), despite reduced phase precession in comparison to neuron pairs from control rats (Fig. 6h, right) and unlike what has been previously reported in CA1 (Schlesiger et al., 2015). Thus, it seems that though the MEC is critical in maintaining hippocampal phase precession, the recurrent CA3 network together with its reciprocal connections with the DG is sufficient to organize the fine ensemble-level temporal relationships of CA3 spiking within theta cycles that compresses behavioral-scale sequences to time-scales suitable for spike-time dependent plasticity.

**Figure 6.**
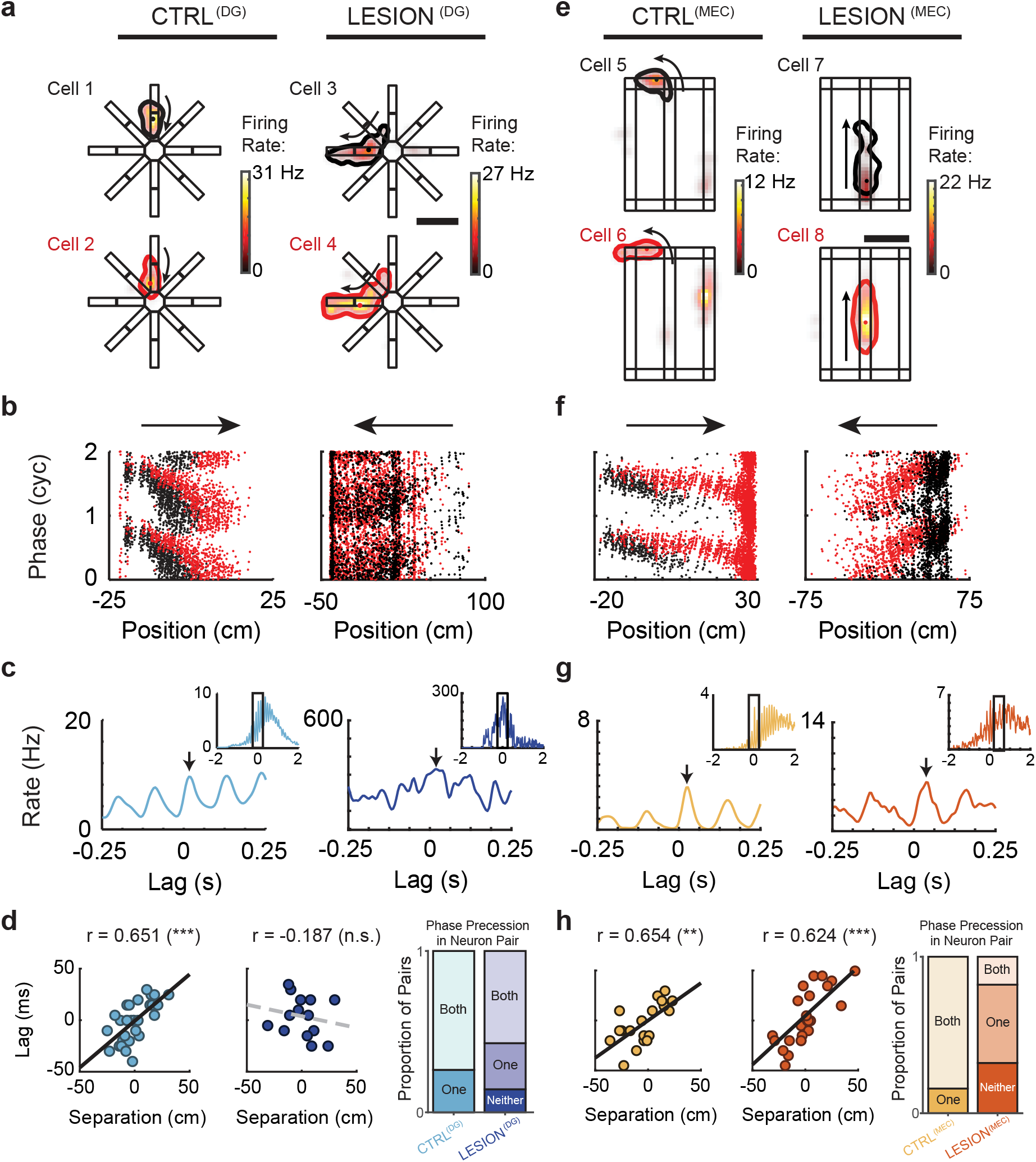
DG but not MEC inputs are required for the temporal organization of CA3 cell pairs at the theta-cycle time scale. **a**, Pairs of simultaneously recorded CA3 cells with overlapping place fields were selected for the analysis of spiking in shared theta cycles. Cells 1 and 2 are from a control rat, cells 3 and 4 from a DG-lesioned rat. Each place field is delineated by a contour that corresponds to 20% of the maximum firing rate (dot inside contour, location of peak firing). Each pair of overlapping fields are depicted with the one that is entered first in black and the one entered second in red. Black arrow, running direction. **b**, Phase-position plots of the cell pairs’ spikes (red and black dots, from the cells depicted in red and black in **a**) while running in the direction indicated by the horizontal arrows (corresponding to the direction in **a**). **c**, Cross-correlation of the spikes (arrow, peak of the cross-correlation function nearest zero lag; inset, cross-correlogram for a window width of 4 seconds). **d**, Left, Phase shift at the theta-cycle time scale plotted against the distance between place field peaks (solid and dashed lines, linear regression for data from control and DG-lesioned rats; CTRL^(DG)^: *n* = 30 pairs, r = 0.651, *p* = 9.87 × 10^−5^, Pearson’s correlation). Right, Proportion of cell pairs where neither, one, or both neurons in the pair displayed phase precession. Although a substantial number of pairs in the DG-lesioned group were phase precessing, the phase precession did not yield a relation between pairwise spike-time difference and distance between fields (LESION^(DG)^: *n* = 14 pairs, r = −0.187, *p* = 0.523, Pearson’s correlation). **e**-**h**, Same as **a**-**d** except that the CA3 data are from the MEC-lesioned group and their respective controls. Despite marked deficits in phase precession, there is a strong correlation between spike-time difference and place field distance (CTRL^(MEC)^ *n* = 19 pairs, r = 0.654, *p* = 0.0024; LESION^(MEC)^ *n* = 27 pairs, r = 0.624, *p* = 5.02 × 10^−4^; Pearson’s correlations). n.s., not significant, ** *p* < 0.01, *** *p* < 0.001.

### A computational model of phase precession in CA3 cells revealed distinct effects of DG and MEC inputs on the inhibitory signal

To further examine how each of the two theta-modulated inputs to CA3 can have distinct effects on phase precession, we devised a minimalistic phenomenological model based on oscillatory interference to simulate the spiking dynamics of a model CA3 pyramidal cell as the animal moves through the cell’s place field at constant velocity. Although our experimental manipulations included two excitatory inputs to CA3, we reasoned that if a computational model based on oscillatory interference were to emulate the lesions, it must account for inputs beyond the manipulated inputs. Inhibition has been shown to mediate input gain control, precise spike timing and enhanced coding in networks (Lytton and Sejnowski, 1991; Klyachko and Stevens, 2006; Milstein et al., 2015; Denève and Machens, 2016) and can thus be considered essential for controlling the theta phase of pyramidal cell spikes (Losonczy et al., 2010). Therefore, we included an inhibitory oscillation in the model that can be viewed as corresponding to the observed oscillations of a large fraction of hippocampal interneurons at the LFP theta frequency (Csicsvari et al., 1999; Klausberger et al., 2003). Based on recordings from DG and MEC principal cells that are known to project to CA3, the excitatory inputs from each of these two regions were considered to oscillate at frequencies slightly above the LFP theta frequency (Jeewajee et al., 2008; Mizuseki et al., 2009). In addition, the relative contributions of DG and MEC inputs varied along the place field to reflect the proposal that entorhinal inputs provide sensory cues at the true place field location while intrahippocampal circuits govern the prospective spiking (Jensen and Lisman, 1996; Mehta et al., 1997; Lisman et al., 2005; Sanders et al., 2015). We estimated and fixed the phase offset between the two oscillations that modeled DG and MEC inputs at ψ = −39° based on data from Mizuseki et al. (2009; their Figure 3). We allowed the model to have three free parameters: a phase shift between excitation and inhibition denoted by ϕ_inh_, the oscillatory amplitude of inhibition denoted by A, and a DC component for the inhibitory oscillation (baseline inhibition) denoted by I_DC_. The output of the model CA3 cell was determined by the place modulated (Silver, 2010; Schmidt-Hieber and Nolan, 2017) combination of the three inputs and from which a threshold value that was constant across the place field was subtracted. Spikes were generated stochastically via an inhomogeneous Poisson point process with an intensity measure defined by the total excitatory drive minus the threshold. The simulated spike phases were extracted with respect to an 8 Hz oscillation representing the LFP theta oscillation, which was considered to be in sync with the inhibitory oscillation (see Methods).

We simulated CA3 model neuron spikes for a broad range of parameter values. We observed for the full model (Fig. 7a) that phase precession can be obtained for a wide range of parameter values (Fig. 7b), but was reduced or non-existent depending on the particular combinations of A and ϕ_inh_ when either of the two excitatory components was removed (examples shown in Fig. 7c). To determine how parameter values under this model corresponded to the experimental data from the control and lesion groups, we calculated four phase precession measures – slope, onset phase, offset phase, or explained variance r^2^ – from the model spike data in the full A versus ϕ_inh_ parameter space. We then identified the region of the parameter space in which the model generated phase precession measures that corresponded to those obtained in our empirical data (i.e., the middle two quartiles of the empirical observations in the experiments). This was done separately by matching the control data to the control model and the lesion data to each of the respective lesion models. In the full control model, only a small range of phase shifts (ϕ_inh_) between the inhibitory input and the excitatory inputs in the model generated phase precession measures that corresponded to data from control animals. When the DG input was set to zero in the model, the model-generated phase precession data matched with the empirical data over a broader and shifted set of ϕ_inh_ values compared to controls (Fig. 7d,e; effect size of comparison between lesion and full model: A: *d* = 0.27, ϕ_inh_: *d* = −2.89). This suggests that DG inputs restrict CA3 spiking activity in the initial part of a place field to a reduced inhibition-mediated later phase window. In contrast, when MEC-lesioned empirical data were matched with model data in which the MEC input was set to zero, the model data that could reproduce the empirical data included values of A covering a broader range and exhibiting a lower mean as compared to the control empirical/model data match (Fig. 7f,g; effect sizes of lesion in comparison to full model: A: −2.0, ϕ_inh_: 0.60). This suggests that the addition of MEC inputs requires larger inhibitory amplitudes and that increased phase variability observed in MEC-lesion data might result from generally reduced theta-modulated inhibition. Interestingly, the third free parameter in the model – the DC component – did not influence the state space substantially as its addition to the model did not produce substantially distinct ranges of match with empirical data (Fig. S5). Since the difference between the match of lesion data to the model primarily arises along two independent dimensions (ϕ_inh_ and A for DG and MEC, respectively), we found it plausible that the inhibition phase compared to the excitatory inputs is mostly set by the DG inputs, whereas the magnitude of the inhibition amplitude is mostly set by the MEC inputs. Taken together, these simulations demonstrate that the range of observed effects in the CA3 circuit can, in principle, be generated by the interaction of the two major theta-modulated excitatory inputs with the local inhibitory neuronal population connecting to CA3 pyramidal cells.

**Figure 7.**
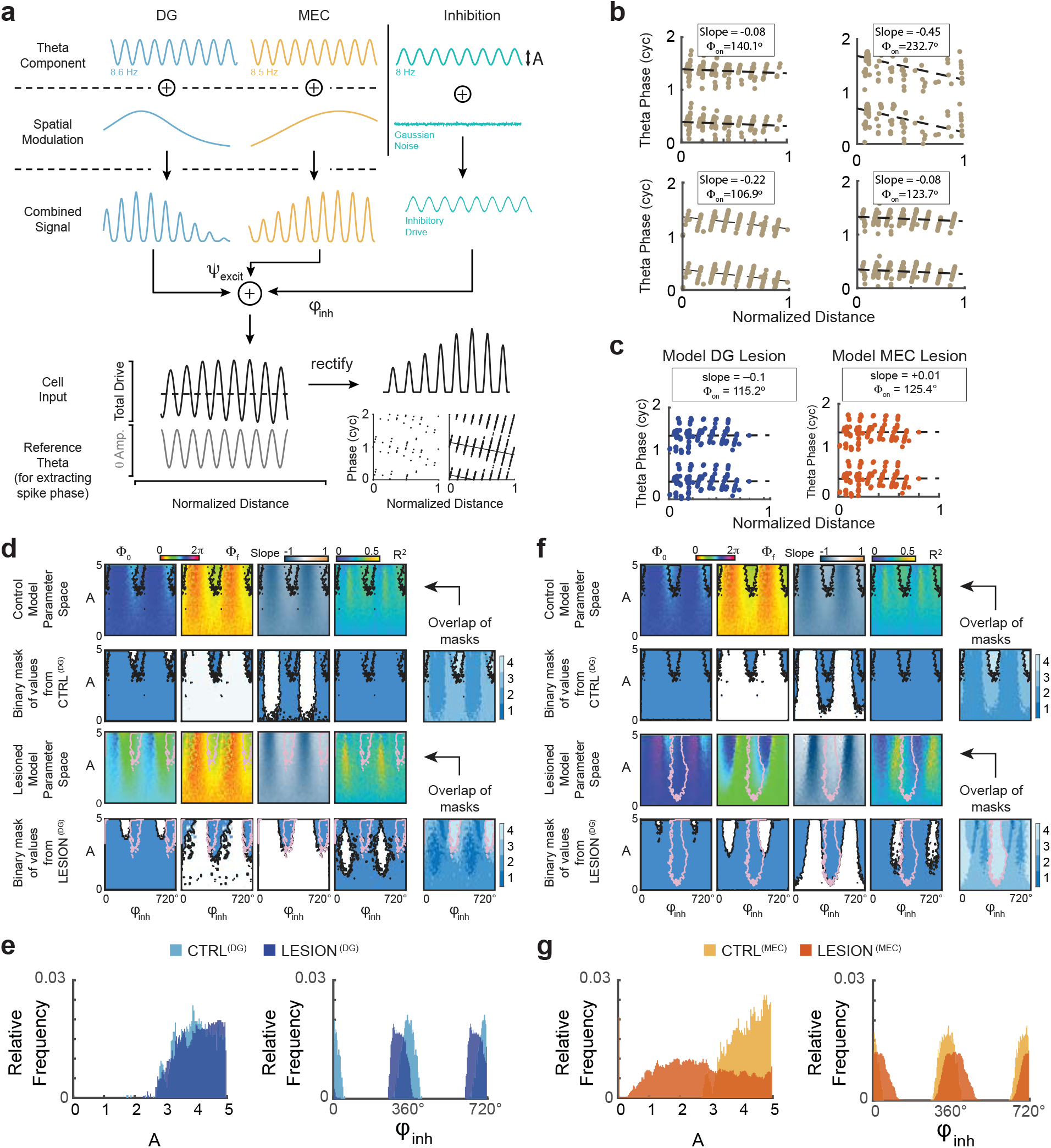
A model of two oscillating excitatory inputs and an inhibitory input reproduced the main empirical results. **a**, Model construction. The three inputs are modeled after DG, MEC, and local inhibition converging onto the CA3 model neuron. The excitatory DG and MEC inputs oscillate at faster-than-LFP frequencies (ω_DG_ = 8.6 Hz, ω_MEC_ = 8.5 Hz) with DG inputs more prominent early in the field and MEC inputs more prominent later in the field. The inhibitory input oscillates at 8 Hz throughout the place field, corresponding to LFP theta. Small Gaussian noise is added to the inhibitory input to ensure robustness against minor perturbations. The excitatory inputs contribute positively at the fixed phase differential ψ, which is taken from published findings (Mizuseki et al., 2009) on DG and MEC population activity. The inhibitory input contributes negatively to the total drive at a phase differential ϕ_inh_ relative to excitation at place field entry. Finally, the total drive is rectified. A reference 8-Hz oscillatory inhibition is displayed at the bottom left, which is used to extract the phase of the simulated spikes. The phase-distance relationship is then depicted as for experimental data. Not all steps are displayed for brevity (see Methods for full details). **b**, Phase-distance relationship of spikes generated by the model show phase precession. The values of A = 1 (inhibitory oscillation amplitude), I_DC_ = 0 (inhibition DC component) and ψ = 158° (excitatory phase differential) are the same across the four plots. However, the inhibition phase offset, ϕ_inh_, varies across panels (clockwise from top left: 0°, 118°, 239°, 299.5°). The measured slope and onset phase of the simulated phase precession are displayed at the top of each subpanel. A broad range of phase precession profiles can be obtained, confirming the expressive power of the simple model. **c**, Lesion experiments were simulated by setting the DG input (left) or MEC input (right) to 0. Values of A = 1, ϕ_inh_ = 118° were used for both panels. The measured slope and onset phase values are displayed on top. In both cases phase precession is diminished, and the slopes are reduced. (**d-g**) DG and MEC lesions alter CA3 phase precession in qualitatively different ways. **d**, Values of onset phase, offset phase, slope, and explained variance (colored plots, I_DC_ is fixed at 0) with various values of A (inhibitory amplitude) and ϕ_inh_ (inhibitory phase shift), and the overlap between model and empirical values (blue and white plots, with blue depicting the area of overlap). The intersection where multiple empirical and model measures overlap is displayed in the overlap plots to the right (dark to light blue, 1 to 4 measures overlap). The overlap of all four measures is delineated with a black (control) or pink (lesion) outline, and the zone of overlap is projected back onto the model parameter space. **e,** Distribution of parameter values A and ϕ_inh_ that result in a match with the empirical control (light blue) and lesion (dark blue) data from the DG lesion experiment (effect size of differences in parameters to fit empirical data to either the lesion or the full model: A: *d* = 0.27, ϕ_inh_: *d* = −2.89). The values in the histograms correspond to those inside the black and pink outlines in **d**. To match the empirical phase precession measures in the DG-lesioned compared to control rats, the model is forced to take on a shifted set of ϕ_inh_ values. **f**, Similar to **d**, but showing the results of lesioning the MEC input in the model. **g**, As in **e** but for the MEC lesion experiment. In contrast to the DG lesion model, the MEC lesion model is forced to take on a broader set of A values to match the empirical phase precession measures (also compare black and pink outlines in **f**; effect size of differences in parameters to fit empirical data to either the lesion or the full model: A: −2.0, ϕ_inh_: 0.60).

## Discussion

The DG is the first processing stage in the hippocampal circuit and is considered to perform a number of specialized computations that are critical for memory such as spatial and temporal pattern separation as well as novelty detection. Furthermore, computational models also emphasize that the dentate-CA3 network forms a loop that could be used for generating and storing sequences (Lisman et al., 2005), which in turn can be used for guiding ongoing behavior and decisions (Gupta et al., 2010; Pfeiffer and Foster, 2013; Singer et al., 2013; Joo and Frank, 2018). While there is recent evidence for a contribution of DG to the activation of CA3 ensembles during SWRs (Sasaki et al., 2018), the role of DG inputs to CA3 during periods when theta oscillations are predominant has not been established. Here we show that diminished DG inputs to CA3 cells resulted in a substantial disruption of precise spike timing within theta cycles and in reduced theta phase precession. The reduced phase precession was accompanied by a disrupted temporal order of the spiking within a theta cycle for cells with overlapping place fields. It is possible that effects on the temporal activity patterns of CA3 cells are not specific to DG inputs but might emerge when any of the theta-modulated excitatory inputs to CA3 are diminished. We therefore compared the effects from reduced DG inputs to CA3 with the effects of reducing MEC inputs. Similar to our observation with DG lesions, we found that loss of MEC inputs resulted in reduced phase precession. Despite the phenomenological similarity when considering standard phase precession measurements, we identified profound differences in the effects of each manipulation on precise timing. Only DG but not MEC lesions precluded spikes from selectively occurring late in the theta cycle at the onset of spike trains, and only DG but not MEC lesions disrupted the pairwise timing between cells that are co-active. Given that manipulations of each of the two theta-modulated inputs to CA3 resulted in distinct effects of spike timing at the theta scale, we generated a model that identified distinct coupling of each of the excitatory inputs to local inhibition as a plausible mechanism of how these differences emerge. By comparing the model to empirical data, we recognized that the effects that resemble DG lesions were more readily achieved by varying the phase of the inhibition while the effects that resemble MEC lesions were more readily achieved by varying the amplitude of the inhibition. Taken together, DG inputs to CA3 therefore have a particularly pronounced role in generating the preferential spiking of CA3 cells during late theta phases and for the temporal order of spiking within a theta cycle.

While standard measurements of phase precession can broadly indicate that spike timing is altered, the more detailed measurements in our study provide further insight into the pattern of disruption. The selective effect on late spiking during the initial theta cycles is evident in the finding that diminished DG inputs preferentially broadened the phase of spiking within a theta cycle at the onset of a spike train, and to a lesser extent, at the offset. We note that these phase shifts were unlikely a result of traveling wave theta phase differences due to tetrode placement (Muller et al., 2018) as this would have altered onset and offset distributions to the same extent, contrary to our observations. Rather, in-depth analyses of spiking during theta cycles revealed that DG lesions resulted in a broadening of the phase window during which spikes are generated during the initial theta cycles. In addition, we also found that the relative timing of CA3 cell pairs on a theta-cycle time scale depended on DG. Importantly, neither the pronounced broadening of the onset phase nor the selective effects on spike timing at the onset of spiking were observed with MEC lesions, which nonetheless reduced phase precession in CA3 to a similar extent as the DG lesions. However, the preserved pairwise spike timing of CA3 after MEC lesions and the moderately preserved phase precession in CA3 after MEC lesions differ from the previously reported profound disruption of the temporal order in pairs of CA1 cells and of phase precession in CA1 cells (Schlesiger et al., 2015; Chenani et al., 2019). The less pronounced effect on the timing of CA3 firing patterns compared to CA1 firing patterns with MEC lesions could be a consequence of the additional inputs from DG to only CA3 but not CA1. These inputs confer the CA3 circuit with the propensity to generate sequential activity patterns, such that this computation – when MEC inputs are diminished – can emerge with remaining DG projections to CA3 but not with remaining CA3 projections to CA1 (Schlesiger et al., 2015). The recurrent CA3 network is thus by itself not sufficient for sequence generation, but as our data suggests, requires the broader DG-CA3 circuit to support these computations.

How is the DG-CA3 projection specialized to support the emergence of precise spike timing? Initial models of the DG contribution to phase precession have emphasized the strong facilitation at mossy fiber synapses from DG granule cells to CA3 pyramidal cells (Thurley et al., 2008). In this scenario, a weak excitatory input would initially not overcome inhibition until facilitation has occurred by repeated activation of the synapse across theta cycles, which would result in increasing facilitation such that inhibition is exceeded at progressively earlier theta phases across theta cycles. However, this straightforward model is not consistent with more recent data, which show that inputs from granule cells to inhibitory interneurons in CA3 will result in feedforward inhibition that at least matches, if not exceeds, the facilitation at mossy fiber synapses to CA3 pyramidal cells, in particular at the time scale across theta cycles (Zucca et al., 2017; Neubrandt et al., 2018), which is particularly relevant to phase precession.

Given the major role of inhibition in shaping the spike timing in intact neural circuits, we asked by introducing a phenomenological model whether our empirical observations could be explained by effects on inhibition. The phenomenological model does not only consider DG and MEC inputs to CA3 but also asks whether the modulation of rhythmic inhibition governs the dynamics of CA3 place field firing. Accordingly, our model CA3 pyramidal cells received the two excitatory inputs from MEC and DG as well as an oscillatory inhibition. The model is therefore conceptually related to previous models of phase precession that have considered oscillatory interference between an inhibitory somatic drive and dendritic excitation (O’Keefe and Recce, 1993; Kamondi et al., 1998; Losonczy et al., 2010) except that two independent excitatory inputs were used here. Model parameters included the theta phase shift between the excitatory and inhibitory inputs and the oscillatory amplitude of inhibition. By exhaustively searching the parameter space of the model, we identified model parameters that gave rise to phase precession measures that corresponded to the empirical observations. We then eliminated either the DG or the MEC input to the model and repeated the search for a feature space that corresponded to the empirical data. By mapping the experimental findings to the computational model, we found that the results point to the DG and MEC input pathways exerting two types of effects on their associated inhibitory subnetworks. The consequences of loss of DG inputs can be explained by a shift in the phase of inhibition, whereas the consequences of loss of MEC inputs can be explained by a decrease in the amplitude of the inhibitory theta signal. Interestingly, this result is consistent with fast-spiking interneurons being targeted by granule cell projections to CA3 (Szabadics and Soltesz, 2009; Royer et al., 2012). In contrast, MEC inputs are known to not only target pyramidal cells but also somatostatin interneurons that predominantly control dendritic inhibition, and manipulations of dendritic inhibition have been shown to be without effect on the average spike phase throughout the place field (Royer et al., 2012), which resembles our observation that MEC lesions do not alter the mean phase. Together, these data therefore suggest that effects from manipulating excitatory inputs do not only arise from diminished direct connectivity to principal cells, but also from how these inputs engage inhibitory interneurons. In particular, our data are consistent with the mossy fiber inputs to CA3 more strongly engaging somatic inhibition, which determines the theta phase of spikes, and with MEC more strongly engaging dendritic inhibition which does not directly set the theta phase.

It is generally assumed that the precisely timed theta phase is a prerequisite for generating sequences of neuronal firing patterns (Foster and Wilson, 2007; Feng et al., 2015), but the circuits that are necessary for implementing these computations have not been established. Because of the much more pronounced effects of DG lesions compared to MEC lesions on the emergence of temporally ordered timing within the theta cycle, our data are consistent with computational models that do not only require the recurrent loops within CA3 for generating sequential firing patterns, but also the loop from the dentate gyrus to CA3 and back (Hasselmo et al., 2002; Lisman et al., 2005; Sanders et al., 2015). Furthermore, our data identify that it is in particular the spike timing in the initial theta cycles that requires DG input. These are the spikes that are thought to emerge from internally generated sequences and argue for a role of DG for biasing CA3 neuronal activity towards such prospective neuronal activity, which goes beyond the previously established functions of DG for pattern separation and novelty detection (Treves and Rolls, 1992), but resembles our previous report that the DG network contributes to prospective neuronal activity patterns during SWRs (Sasaki et al., 2018). Here we show that emergence of ‘look-ahead’ spiking during theta states also depends on DG inputs to CA3, which broadly implicates the DG in sequence coding and future planning, in addition to its well-established functions in pattern separation and novelty detection.

## Acknowledgements

We thank M. Wong and A.-L. Schlenner for technical assistance. This work was supported by NIH grant R01 MH119179 and the Walter F. Heiligenberg Professorship to J.K. Leutgeb, NIH grants R01 NS102915, R21 MH100354, R01 NS084324, and R01 NS097772 to S. Leutgeb, and DFG (German Research Association) grant LE2250/13-1 to C. Leibold. M. Sabariego. was supported by NIH training grant T32 AG 00216.

## Author Contributions

S.A., S.L. and J.K.L. conceived experiments, designed study, and interpreted data. S.L. and J.K.L. managed the project. T.S. and M.S. prepared and provided data from previously published experimental work, S.A. analyzed data, S.A. created computational model in collaboration with C.L. S.A. and J.K.L. prepared figures, and S.A., S.L., and J.K.L. wrote the manuscript with feedback and editing from C.L.

## Methods

### Subjects and Surgical Procedures

All experimental procedures can be found in previous publications describing data analyzed in the present study (Sasaki et al., 2018; Sabariego et al., 2019). We reanalyzed and compared CA3 activity patterns from these data, including a total of 31 rats (Table S1). These included 4 control and 9 dentate-lesioned rats (DG lesion experiment) with CA3 or dual CA3-DG (2 of the 4 control rats) single-unit recordings, 7 control and 8 MEC-lesioned rats (MEC lesion experiment) with CA3 single-unit recordings, and 3 control rats with only DG recordings. Male Long-Evans rats between the ages of 3 and 6 months (300-350g) were used as subjects. The animals were kept on a 12-hour light-dark cycle (7 AM to 7 PM dark) and housed individually. In vivo recordings were conducted in the dark phase. Rats were restricted to 85% of their ad libitum weight and given full access to water. All procedures were conducted in accordance with the University of California, San Diego Institutional Animal Care and Use Committee, at the University of California, San Diego according to National Institutes of Health guidelines.

### Experimental Procedures and Brain Lesions

The details of the DG lesion and MEC lesion experiments were published previously (Sasaki et al., 2018; Sabariego et al., 2019). In brief, rats in the DG lesion experiment were trained on the 8-arm radial maze to perform a spatial-working memory task (see Behavioral Tasks section). Rats initially designated to receive DG lesions (*n* = 9 animals; LESION^(DG)^) underwent a surgical procedure during which colchicine was bilaterally infused along the septal-temporal axis of the DG. The remaining rats (*n* = 4 animals; CTRL^(DG)^) were subjected to a sham lesion. During the same surgical procedure, a hyperdrive of 14 independently moving tetrodes was implanted above the right hippocampus for electrophysiology as described below. Rats in the MEC lesion experiment were trained on a working memory task, the figure-8 continuous spatial alternation task (see Behavioral Tasks section). These rats underwent a surgical procedures in which the control rats (*n* = 7 animals; CTRL^(MEC)^) received a sham lesion (injection of vehicle) and the experimental rats (*n* = 8 animals; LESION^(MEC)^) received an excitotoxic lesion of MEC by an injection of NMDA.

In post-mortem histological material, the final position of the recording tetrodes was confirmed by performing cresyl violet staining of the sectioned brain tissue. In the DG Lesion experiment, the loss of dentate granule input to CA3 cells was confirmed by TIMM stains as previously described in detail (Sasaki et al., 2018). The extent of DG granule cell damage was quantified in a localized fashion. Specifically, each tetrode ending location was scored based on the intensity of the TIMM-positive staining in histological sections. Scores of 0 (∼0% TIMM-positive signal), 1 (< 30% signal), or 2 (< 70% signal), 3 (> 70% signal) were assigned to each of the tetrodes in DG lesioned animals, and only tetrodes with scores of 0, 1, and 2 were included in the LESION^(DG)^ data set. The extent of MEC lesions was confirmed quantitatively (Sabariego et al., 2019), and on average, 93.0% of the total MEC volume was ablated (95.3% of layer II, 92.4% of layer III, and 91.4% of deep layers) with any sparing typically observed in the most ventral portions of MEC.

### Hyperdrive Implants

An array of 14 independently movable tetrodes was implanted over the right hippocampus in all 31 rats (control group: 4.0 mm posterior and +2.7 to +2.9 mm lateral to bregma; DG lesion group: 3.5 to 4.4 mm posterior and +2.8 to +3.2 lateral to bregma; MEC lesion group: −4.0 mm posterior and +2.8 lateral to bregma). The hyperdrive was secured with skull screws and dental cement to prevent mechanical instability. The tetrodes (with tips platinum plated to 150-300 kΩ at 1 kHz) were slowly lowered each day over a period of 2-4 weeks to ensure recording stability and minimizing damage to the brain. Depth records, LFP signals, and neural spiking markers were used to estimate tetrode distance from the target region. After an initial period of larger advances, the tetrodes were moved only in small increments over several days until a satisfactory signal (i.e., low-amplitude multiunit activity) was observed. Once near CA3, the tetrodes were allowed to settle inside the stratum pyramidale of the CA3 of the hippocampus with no further active movement of the tetrodes to maximize recording quality (i.e., high-amplitude multiunit activity). In 3 of the 31 rats all tetrodes were lowered to the dentate granule layer.

### Electrophysiological Recordings

A Neuralynx Cheetah recording system with a multichannel head-mounted preamplifier was used for LFP and single-unit recordings. A signal from a skull screw was used as animal ground, and a reference signal from the neocortex was subtracted from the hippocampal signals to increase the hippocampal signal to noise ratio. Unit recordings were filtered at 600 Hz to 6 kHz, and spike waveforms above an amplitude of 40 μV were time-stamped and recorded at 32 kHz for 1 ms. LFP recordings were filtered between 1 and 425 Hz in the DG lesion experiment and between 1 and 450 Hz in the MEC lesion experiment.

### Behavioral Tasks

#### DG Lesion Experiment (Spatial working memory on the 8-arm radial maze)

The rats in the DG lesion experiment were trained to perform a DG-dependent spatial working memory task (Sasaki et al., 2018). The task used a maze with a central platform and 8 radial arms that each had a proximal segment that could be lowered and raised. The rats were first placed on the central platform (i.e., “stem”) of the 8-arm maze with all 8 arms lowered such that the reward cups at the end of each arm were inaccessible to the animal (Fig. S1a). Next, the experimenter raised one arm at a time, for four arms, following a previously generated pseudorandom sequence. The rat was allowed to run down each raised arm and upon its return to the stem that arm was lowered and the next arm in the sequence was raised. Once the rat had visited all four experimenter-forced arms (“forced choice” phase), all 8 arms were raised and available for the rat to visit (“free choice” phase). The optimal strategy would consist of the rat visiting every one of the four arms unvisited during the forced-choice phase without reentering any arm. A total of 16 rats (*n* = 4 CTRL^(DG)^; *n* = 9 LESION^(DG)^, *n* = 3 with only DG recordings) were trained and tested in this task while performing single-unit recordings were acquired in CA3 and/or DG.

#### MEC Lesion Experiment (Spatial working memory on the Figure-8 maze)

In the MEC lesion experiment, the rats were trained to perform a hippocampus dependent alternation task on the figure-8 maze (Sabariego et al., 2019). In this task, a rat is placed in a delay zone at the base of the figure-8 maze (Fig. S1b) and is required to run up the “stem” of the maze toward a T-junction from where it can choose between reward locations on either the left or right before returning on a side arm to the delay zone. Blocks of trials with and without delays were performed. In non-delayed trial blocks the delay site is not used to restrict the animal’s movements. In delayed trial blocks a barrier restricted the rat’s progress for 2, 10, or 60 seconds for each trial. These blocks were not distinguished for the analyses presented here. This first lap (“trial 0”) is discarded. However, it is used to determine the success of the animal in choosing the right or left reward on the following lap. From the second lap onwards (trial 1 and later), a trial is “correct” if and only if the animal chooses to visit the reward location not visited on the preceding trial. If the animal chooses the same side more than once, it will not receive a reward at the visited reward site. It will, however, continue to receive reward items at the appropriate reward sites as soon as it chooses the side not chosen on the previous trial. A total of 15 rats (*n* = 7 CTRL^(MEC)^; *n* = 8 LESION^(MEC)^) were trained and tested on this task while CA3 recordings were performed.

### Data Analyses

All statistical tests were chosen to appropriately match the underlying data distributions. In the case of testing proportions, *χ*^2^ tests were used. In the case of testing for means, first the normality and homoscedasticity were tested with Anderson-Darling and F-test, respectively. If the data were concluded to be normal, t-tests or ANOVAs were performed as appropriate. Otherwise, Wilcoxon rank-sum or Kruskal-Wallis tests were applied as appropriate. For comparing the distributions, the Kolmogorov-Smirnov test was used. The α level was set to 0.05 for all experiments and tests.

#### Spike Sorting

Spike sorting was performed offline using a custom version of MClust 3.5 (Redish, A.D., http://redishlab.neuroscience.umn.edu/MClust/MClust.html). Clusters were selected in the sleep sessions before and after a behavior and matched to the data recorded during the behavior to ensure consistency and reliability. The cross-correlogram was used as an additional criterion to ensure cluster independence. Only well separated clusters were retained for analysis.

#### Spatial Firing Properties

The measures included in Fig. S2 are defined as follows. Let N be the total number of spikes of a given cell, and the number of such spikes that were part of a detected train (as defined below) denoted N_t_. The proportion of spikes assigned to a train is defined N_t_/N. The firing rate is defined as λ = *N*/*T* where T is the summed total duration in seconds of all behavioral trials in a given session. The number of spikes per train is calculated for each cell as the N / N_tr_ where N_tr_ is the number of detected trains for that cell. To calculate the train length we first found the physical position of the first and last spikes of a train on the maze (points P_1_ = (x_1_, y_1_) and P_N_ = (x_N_, y_N_)). The train length L is calculated as 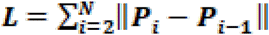, where ||·|| is the L_2_ norm. The number of bins covered was calculated as the total number of square bins of size 2 centimeters that contained at least one point from the path of a detected train. The information content measure was adapted from (Skaggs et al., 1992) and was calculated as 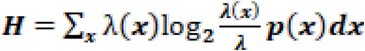, where λ is the mean firing rate and *p* is the probability mass function of the rat’s position over the spatial bins of the maze. The selectivity and sparsity measures were drawn from (Skaggs et al., 1996) and are defined, respectively, as 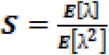 and 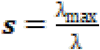, where *E* is the expected value over the spatial bins and λ is the mean rate as defined above.

#### Rate map construction

First, intervals during the periods delimited by trial timestamps in which the velocity of the animal exceeded 2 cm/s were selected. For each cell, all spikes that occurred outside these intervals were excluded for the construction of rate maps. Next, the environment was divided into square bins of side length 5 cm and the spikes that occurred in each such bin were counted. The occupancy matrix was constructed similarly by counting the number of position tracking points falling in each spatial bin multiplied by the tracking acquisition rate (29.97 FPS). The rate map was the result of the element-wise division of the spike count matrix by the occupancy matrix, spatially smoothed with a 2-d Gaussian of kernel size 15 cm.

#### Spike Train Detection

A spike train was defined as the set of 5 or more consecutive spikes with a maximal inter-spike interval of 500 ms. Additional criteria were imposed on the selection of spike trains for analysis (Fig. S2c). A spike train (a.k.a. “pass”) was deemed valid for analysis if it was at least 300 ms and no more than 2500 ms in duration, its corresponding path was at least 20 cm long (see above for the calculation of pass length), its corresponding path endpoints were at least 10 cm apart in physical space, and if the average velocity of the animal during the pass exceeded 2 cm/s. All of a cell’s detected trains were discarded if its mean firing rate over the duration of the behavior was smaller than 0.1 Hz or greater than 5 Hz.

#### LFP Analysis and Theta Phase Extraction

Local field potentials were recorded from one of the electrodes for each tetrode. The raw LFP signal was filtered in the theta range (6-10 Hz) and the channel with the largest theta rectified RMS power was selected as the reference for phase precession analysis. The phase estimate was obtained by 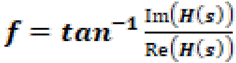 under linear interpolation, where *H*(·) is the Hilbert transform and *s* is the 6 Hz to 10 Hz filtered LFP signal.

#### Quantification of Phase Precession

The distance and theta phase variables were extracted from detected trains. For each train, the last sampled (x,y) coordinate before the first spike and the first sampled (x,y) coordinate after the last spike were marked as the Start and the End of that train’s corresponding trajectory. Next, spike positions were normalized with respect to the Start and End points, yielding vector d of normalized distances. Taken together with the theta phase vector *ϑ* described above, the circular-linear regression was then computed on the (*d*, *ϑ*) pairs for each train or cell, as described below. The circular-linear regression produced a slope value *s* and the estimate of explained variance *r*^2^. The onset phase Φ_on_ was calculated for each train as the circular mean theta phase of the spikes occurring in the first (possibly truncated) theta cycle of the train. The offset phase Φ_off_ was defined similarly, except over the last theta cycle of the train. These four values are referred to as phase precession “measures” in the modeling section.

#### Population level quantification of remaining phase precession

Two methods were employed for the quantification of the phase precession in each data set: slope-by-cell analysis and slope-by-train analysis.

##### Slope-by-cell analysis

In this method, for each CA3 cell, we performed the circular-linear regression analysis on the set of all spikes that belonged to a detected train from that cell. A cell was deemed to exhibit “phase precession”, if the circular-linear regression *p*-value was less than *α* = 0.05 and the slope was negative. The quantification of proportions was performed on the values thus obtained.

##### Slope-by-train analysis

Here, the circular-linear regression was performed on the spikes from individual trains *i* of each cell, to produce phase precession measures. To obtain a cell-specific slope value, the *s*^(*i*)^ values were averaged for each cell. The statistical comparisons of proportions were then performed on the cell-averaged values (such that each cell contributed a single slope value regardless of the number of its trains). The Φ_on_ and Φ_off_ values were only defined for trains, though instead of averaging them per train, they were directly used to find the distributions used in Fig. 3.

#### Statistical tests involving theta phase values

The circular statistics toolbox CircStats (Berens, 2009) was used for the computation of statistical quantities and tests involving values of circular nature, such as theta phase.

#### Spike phase variance analysis

For Fig. 5d, circular variance was calculated either across all spikes of each neuron or across the timestamps resulting from replacing each cycle’s spikes with their mean. Notice that even though this operation reduces the total number of spikes, it will not necessarily reduce the variance; this would depend on the distribution of spikes within the theta cycle and the phase reliability firing windows of a cell over multiple trains. This in turn depends on the brain circuitry which is manipulated by experimental conditions (CTRL, DG lesions, or MEC lesions). For Fig. S3c, we used a similar binning scheme as for Fig. 5d but the binning was performed on the actual place field of the neuron, defined as the area within the 20% contour of the place maps. We then calculated the circular standard deviation across all spikes of each cell in each bin and plotted the mean values together with the error bars representing the standard error of the mean.

#### Onset, offset and binned theta phase estimation

The onset firing phase was defined as the circular mean phase of the spikes occurring in the first (partial) theta cycle of each train. The offset firing phase was analogously defined as the circular mean phase of the spikes occurring in the last (partial) theta cycle of each train. The histograms in Fig. 3 are obtained from the single train onset and offset phase values for each group. To estimate firing probability in the binned theta cycle (Fig. 4), we assigned a bin label (early, mid, or late, corresponding to [0, 2π/3), [2π/3, 4π/3), (4π/3, 2π), respectively) to each spike and plotted the resulting discrete probability distribution (panels a and b). This approach was repeated for each of 10 equal bins of the normalized position of spikes within a train to get a “position-resolved” theta bin firing probability estimate (panel c).

#### Test for common mean in circular data (Circular MANOVA)

As suggested by Landler et al., (2021), we used a one-factor MANOVA test to test for differences in the mean of circular data (e.g., theta phase). Each phase θ was treated as a single observation with two response variables cos(θ) and sin(θ), with the experimental group (control or lesion) as the sole factor. The p-value was automatically obtained by Matlab’s ‘`manova1`’ function by comparing the test statistics with the chi-square distribution with 2 degrees of freedom. This procedure is referred to in the main text as “circular MANOVA.” We avoided using the Watson-Williams test due to its inapplicability when the mean resultant length of the pooled data is < 0.45, as well as the superior performance of MANOVA (Landler et al., 2021). Other tests (permutation tests, Kuiper test, Watson’s U^2^, CircStats (Berens, 2009) test for common medians) returned similar patterns of results (not shown).

#### Test for common concentration parameter in circular data

As suggested by Landler et al. (2021), we used the concentration test from the Directional toolbox (Tsagris et al., 2022). However, we translated this code to Matlab before applying it to data.

#### Calculation of effect size

To quantify the effect size of linear ratio scale data (linear data), we used Cohen’s d defined as 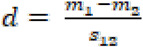 where 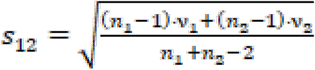, *n_i_* are the number of data points and *v_i_* are the data variances. To quantify effect size for circular data, we used the same formula but using the circular version of mean and variance functions.

#### Cell pair sequence analysis

Two simultaneously recorded units were considered a “pair” for the purpose of sequence analyses if a) there were at least 50 theta cycles in which both units spiked, b) at least 20% of each unit’s spikes throughout the session occurred in theta cycles in which the other unit also spiked, and c) at least 10% of all theta cycles in which a unit spiked included spikes from the other unit as well. For each unit pair, the cross-correlogram was computed and the relative time, τ, of the peak closest to the zero time lag was found. The physical separation of the peaks of the two units’ place fields, d, was computed and used to make the tuple (d, τ). In the case of the 8-arm maze where runs in opposite outbound and inbound directions were possible, each unit pair was treated twice—once for the inbound run and once for the outbound. We confirmed that in all cases only one of the two run directions had enough spikes to reliably assess pair co-modulation (i.e., the unreliable direction did not contribute a (d, τ) tuple). Once all (d, τ) tuples were obtained for each experimental group (CTRL^(DG)^, LESION^(DG)^, CTRL^(MEC)^, and LESION^(MEC)^), a linear regression model was fit to the data to assess the significance of the relationship between place field separation (d) and theta co-modulation (τ), as reported in Fig. 6. For the phase precession plots in Fig. 6b and Fig. 6f, trains from each cell were first mapped to the animal’s path. The path equivalent of each train was then projected onto the line segment connecting the place field peaks of the two cells via a dot product. The midpoint of this line segment was considered the origin (x = 0) and was used on the x-axis of the phase-position plots. The sign of the direction of travel was defined to be positive if Cell 1’s place field was visited before Cell 2’s place field; otherwise, it was considered negative.

### Computational model

#### Model description

The model CA3 neuron received three distinct inputs. Two of these were excitatory (i.e., positively contributed to the model neuron’s total drive) and one was inhibitory (i.e., negatively contributed to the total drive). The excitatory inputs modeled the DG and MEC monosynaptic excitatory drive that CA3 principal cells receive, while the inhibitory input modeled the total inhibitory input that these neurons receive. The equations governing the value of each of the three functions took the following forms:

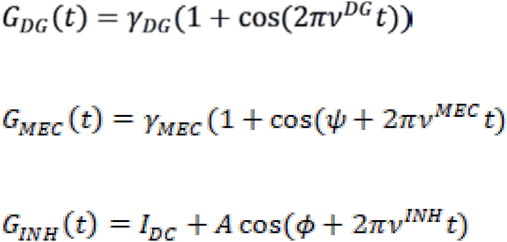

where *ν*^DG^ = 8.6 *Hz*, *ν*^MEC^ = 8.5 *Hz*, and *ν^INH^* = 8 *Hz*, and *I_DC_* represented the DC (baseline) component of inhibition. Gain coefficients *γ_DG_* and *γ_MEC_* were set to 0 to simulate DG and MEC lesion experiments, respectively. In the above equations, *ψ* controls the phase difference of the two excitatory inputs by essentially timing only the MEC input while the DG input remains the same. Effectively, this could cause the place field to shift around slightly which we shall ignore. Here, ϕ denotes the excitatory-inhibitory phase differential, which in the text is referred to by ϕ_inh_ for clarity. I_DC_ was drawn from a Gaussian with a constant mean between 0.5 and 25 (with specific values given in Results) and variance 0.025.

To produce the total drive, the inputs combined as follows:

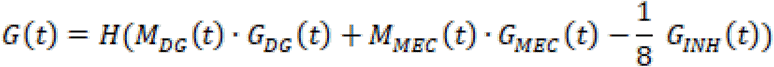

*H* is the Heaviside function (“rectify” step in Fig. 7) and M represents a spatial modulation function. A spatial modulation function to each of these inputs to mimic the influence of DG and MEC inputs to the early and late portions of place fields, respectively (see Fig. 4) (Sanders et al., 2015). These functions are defined as follows:

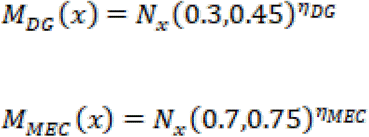

where *η_DG_* = 0.128 and *η_MEC_* = 1.88 are concentration parameters and *N_x_*(*μ*, *σ*) is the Gaussian distribution with mean *μ* and standard deviation *σ*.

#### Spike generation

The total drive obtained in the previous step was normalized to define a probability distribution and used as an intensity function for an inhomogeneous Poisson process to generate the spikes. The phase precession measures were calculated for these simulated trains as described for the empirical data.

#### Mapping of model output to empirical data

The phase precession measures (slope, variance explained, onset phase, offset phase) obtained by simulating the model with various combinations of free parameters (A, I_DC_, and ϕ_inh_) were individually compared to those obtained by analyzing the experimental data from control, DG lesioned, or MEC lesioned rats. Each set of empirical phase precession measures was compared to the phase precession measures obtained from analyzing the corresponding model instantiation (CTRL to full model (no terms set to 0), LESION^(DG)^ to “DG-lesioned” model, and LESION^(MEC)^ to “MEC-lesioned” model). Each measure from the model that was within one quartile (in each direction) of the empirical median of the measures was considered admissible (blue regions in the binary plots of Fig. 7d,e). Free parameter combinations that produced four admissible measures were accepted (Overlap plots in Fig. 7d,e). Finally, the accepted free parameters were compared between lesion and control instantiations of the model by plotting the histogram of their distribution (Fig. 7). The effect size of the change in the admissible parameter spaces was compared by calculating dividing the difference between the mean of control and lesion groups by the pooled standard deviation. The standard deviation was calculated by marking the 2.5% and 97.5% values, taking the difference between the two, and dividing the result by 4 (we assumed the distributions were gaussian, so 95% of the data would lie within 2 standard deviations on either side of the mean).

## Supplemental Information Titles and Legends

**Figure S1.**
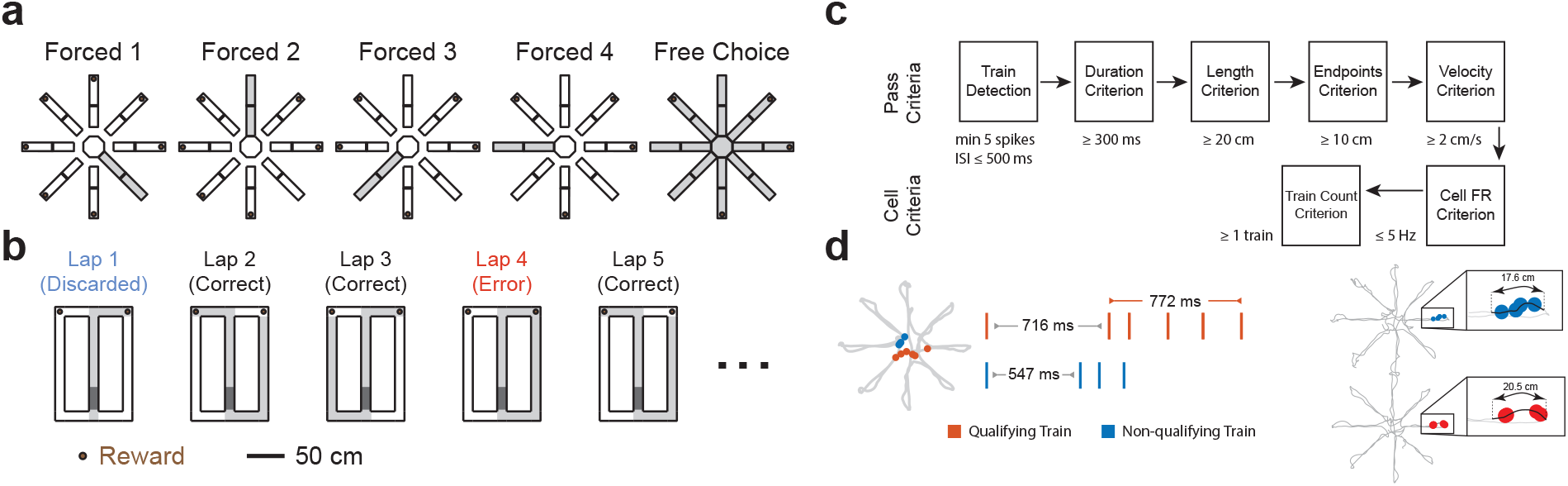
Behavioral tasks and spike train detection. **a**, In the spatial working memory task on the 8-arm radial maze, rats retrieved food from the end of each arm during each trial. The first four arms were chosen in pseudorandom order by the experimenter (Forced 1, Forced 2, Forced 3, Forced 4), and rats then had to collect the food from the remaining four arms (Free Choice). The gray shading indicates accessible arms. **b**, In the delayed alternation task on the figure-8 maze, rats were allowed to choose freely between two reward locations on the first lap and then had to alternate between reward location to receive rewards on subsequent laps after briefly waiting in a delay zone between choices. An example sequence of laps is shown with gray shading indicating the maze regions covered on each lap. The tasks on the radial 8-arm maze task and on the figure-8 maze are both hippocampus-dependent spatial working memory tasks (Ainge et al., 2007; Sasaki et al., 2018). **c**, Spike trains were detected and a series of criteria were then applied to ensure that trains covered a sufficient distance and duration. In addition, cell selection criteria were used to exclude interneurons and to include only cells with a minimum of at least one train. **d**, Examples that qualified or did not qualify as trains. Left, for a CTRL^(DG)^ cell, two spike trains and their corresponding spatial locations (red circles, spikes in qualifying trains; blue circles, spikes that did not qualify for trains due to insufficient number of spikes in train). Right, two trains that qualify based on train detection criteria, but the top example is excluded because the path length during the train is <20 cm.

**Figure S2.**
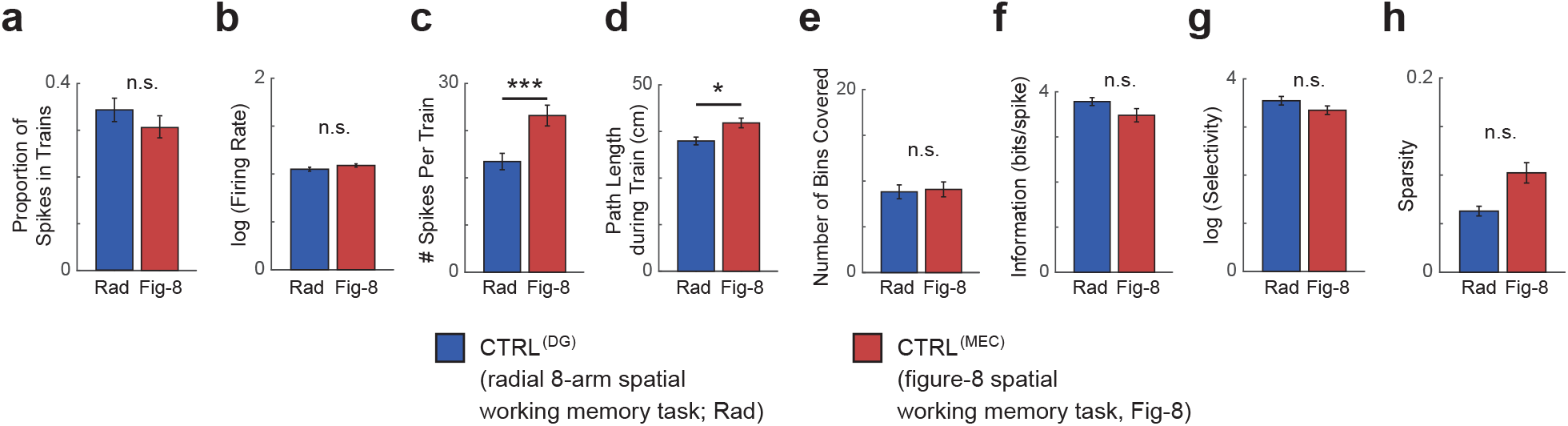
Firing characteristics of trains in control CA3 cells showed only minor differences across the two spatial working memory tasks. **a**, Proportion of recorded spikes that were assigned to trains did not differ between tasks (CTRL^(DG)^ vs. CTRL^(MEC)^, z-statistic = 1.33, *p* = 0.19, Wilcoxon rank sum test). **b-h**, n = 84 and 101 cells for CTRL^(DG)^ vs. CTRL^(MEC)^, respectively. **b**, Average firing rates in identified trains did not differ (log(Firing Rate), log-transformed to obtain normally distributed data, *t*_183_ = −1.47, *p* = 0.14). **c**, Number of spikes per train differed between the two tasks (log-transformed, *t*_183_ = −3.62, *p* = 3.78 × 10^−4^; two-sample *t*-test). **d**, Train length was different across tasks (z-statistic = −2.30, rank sum = 6977, *p* = 0.02, Wilcoxon rank sum test). **e**, The number of bins (2 cm by 2 cm) that contained any portion of a path that was associated with a train did not differ (z-statistic = 0.07, rank sum = 7839, *p* = 0.94, Wilcoxon rank sum test). **f**, Spatial information of spikes in trains did not differ between tasks (z-statistic = 1.33, rank sum = 8295, *p* = 0.18, Wilcoxon rank sum tests). **g**, Selectivity of spikes in trains, as defined by Skaggs et al. (1996), was not different (z-statistic = 1.25, rank sum = 8268, *p* = 0.21, Wilcoxon rank sum tests). **h**, Sparsity of spikes in trains, as defined by Skaggs et al. (1996), was not different (z-statistic = −0.88, rank sum = 7490, *p* = 0.38, Wilcoxon rank sum tests). In addition to confirming that the firing characteristics of trains in control CA3 cells showed only minor differences across the two spatial working memory tasks, potential task-related differences between DG and MEC-lesion effects were further minimized by comparing lesion effects to only the control data from the same task. Bars and error bars, mean ± s.e.m., n.s., not significant, * *p* < 0.05, *** *p* < 0.001.

**Figure S3.**
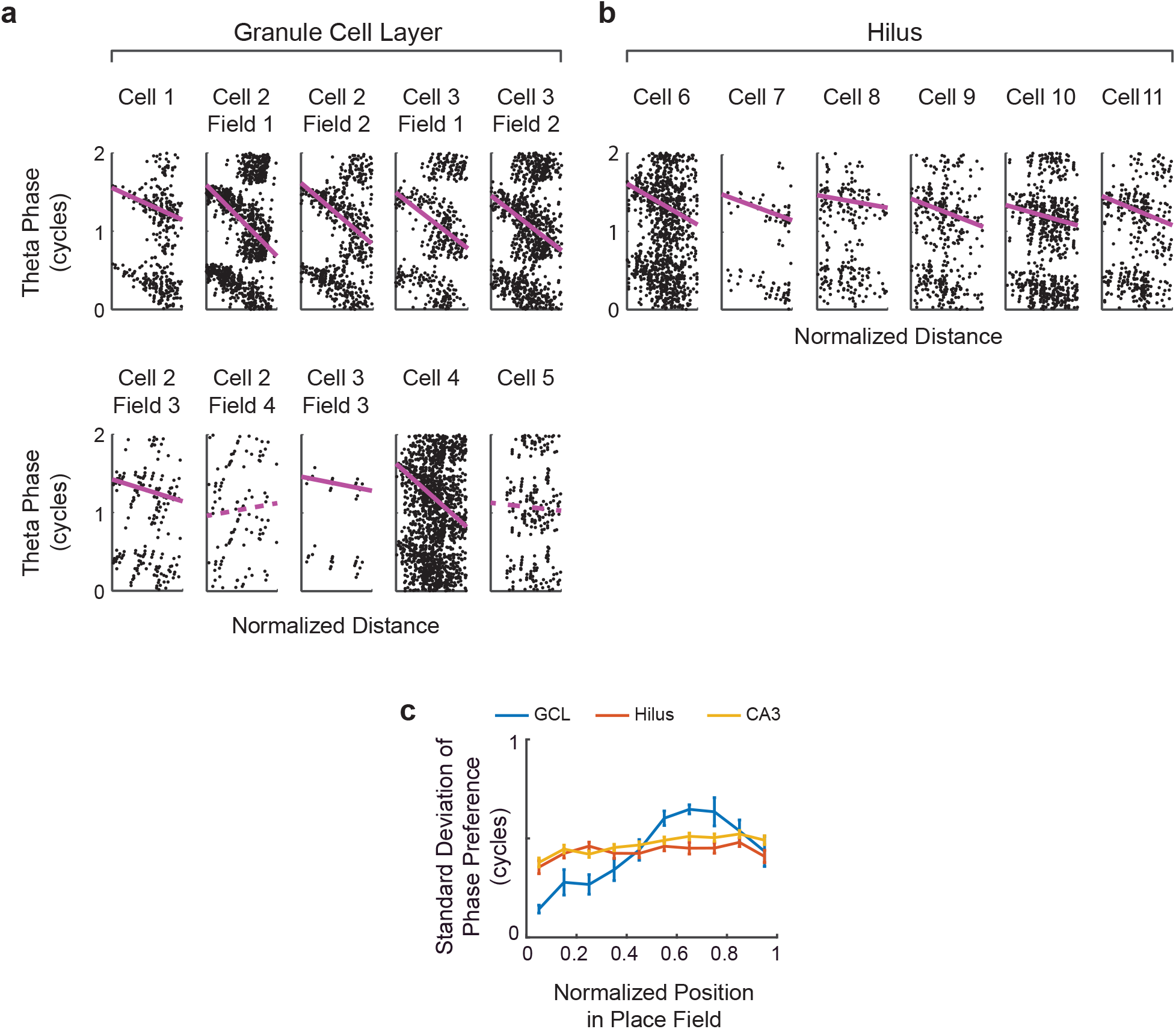
Neurons recorded in the dentate granule cell layer showed the lowest spike phase variability at the entry into place fields. **a**, Phase-distance plots of all neurons that were recorded on tetrodes that were confirmed to terminate in the granule cell layer (GCL, *n* = 5 cells, 10 place fields from 3 rats). **b**, Six phase-distance plots from neurons that were recorded from tetrodes that were confirmed to terminate in the dentate gyrus but in the hilar region or the subgranular zone rather than the granule cell layer (Hilus, total sample, *n* = 13 neurons, 34 place fields from 4 rats). **c**, Quantification of theta phase variability (circular standard deviation) as mean ± s.e.m at various distances through the place field. These plots show that, compared to hilar and CA3 cells, putative granule cells tend to fire at a consistent phase at the entry into the place field. Granule cells could therefore provide a reliable input that controls the onset phase of CA3 pyramidal cell spiking.

**Figure S4.**
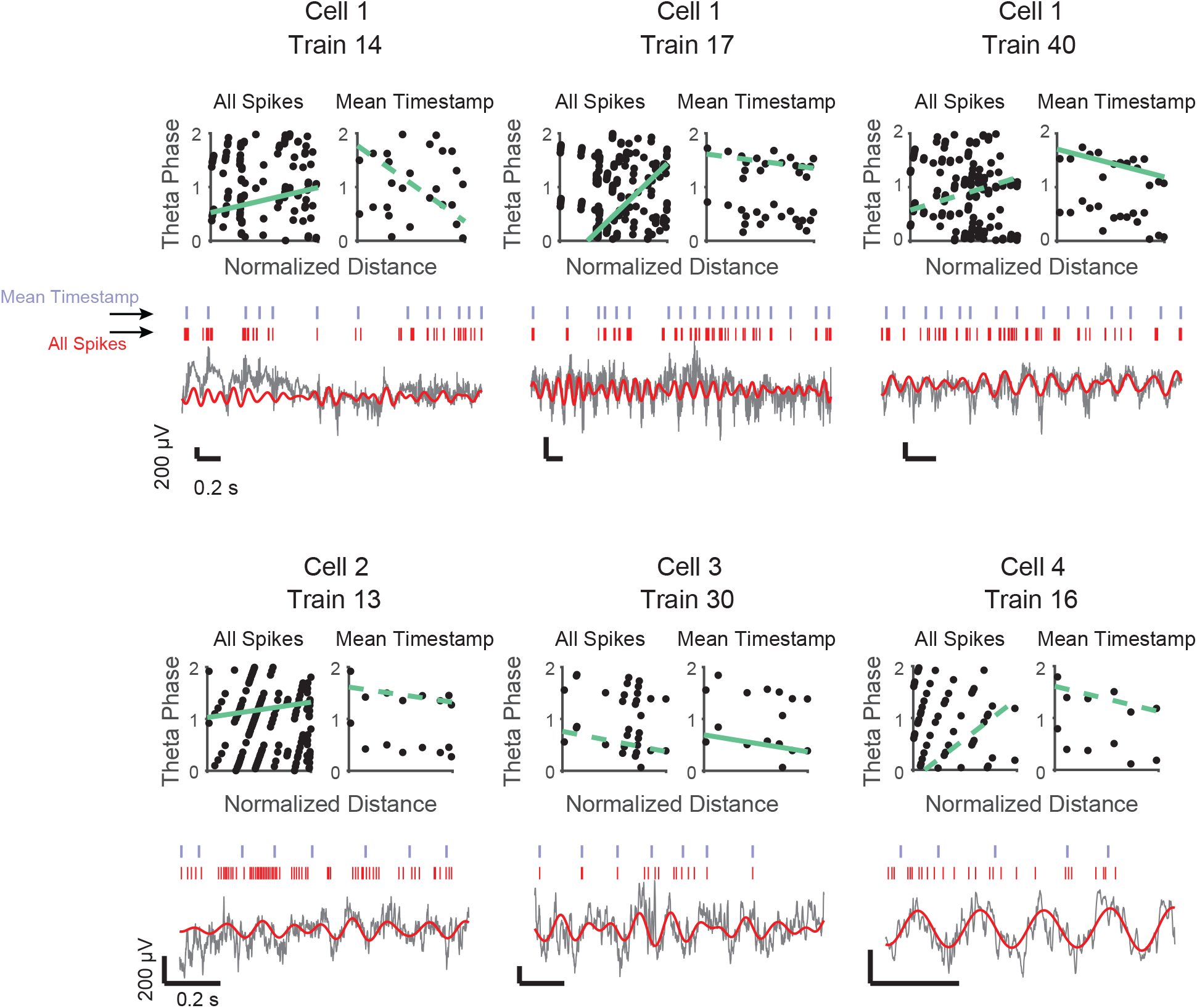
Replacing each theta cycle’s spike phases with the mean phase within the cycle rescued CA3 phase precession in cells from the MEC-lesion, but not from the DG-lesion group. Examples of CA3 single spike trains from the MEC lesion group. Replacing each theta cycle’s spike phases with the mean phase results in a negative slope when a positive slope was detected with the unprocessed spike phases. Below the phase-distance plots, the spike train (red ticks) and the mean timestamps of each cycle’s spikes (blue ticks) are displayed along with the simultaneously recorded LFP (gray, unfiltered, red, 6-10 Hz filtered). Solid green lines, statistically significant single-train slopes, stippled green lines, non-significant single-train slopes.

**Figure S5.**
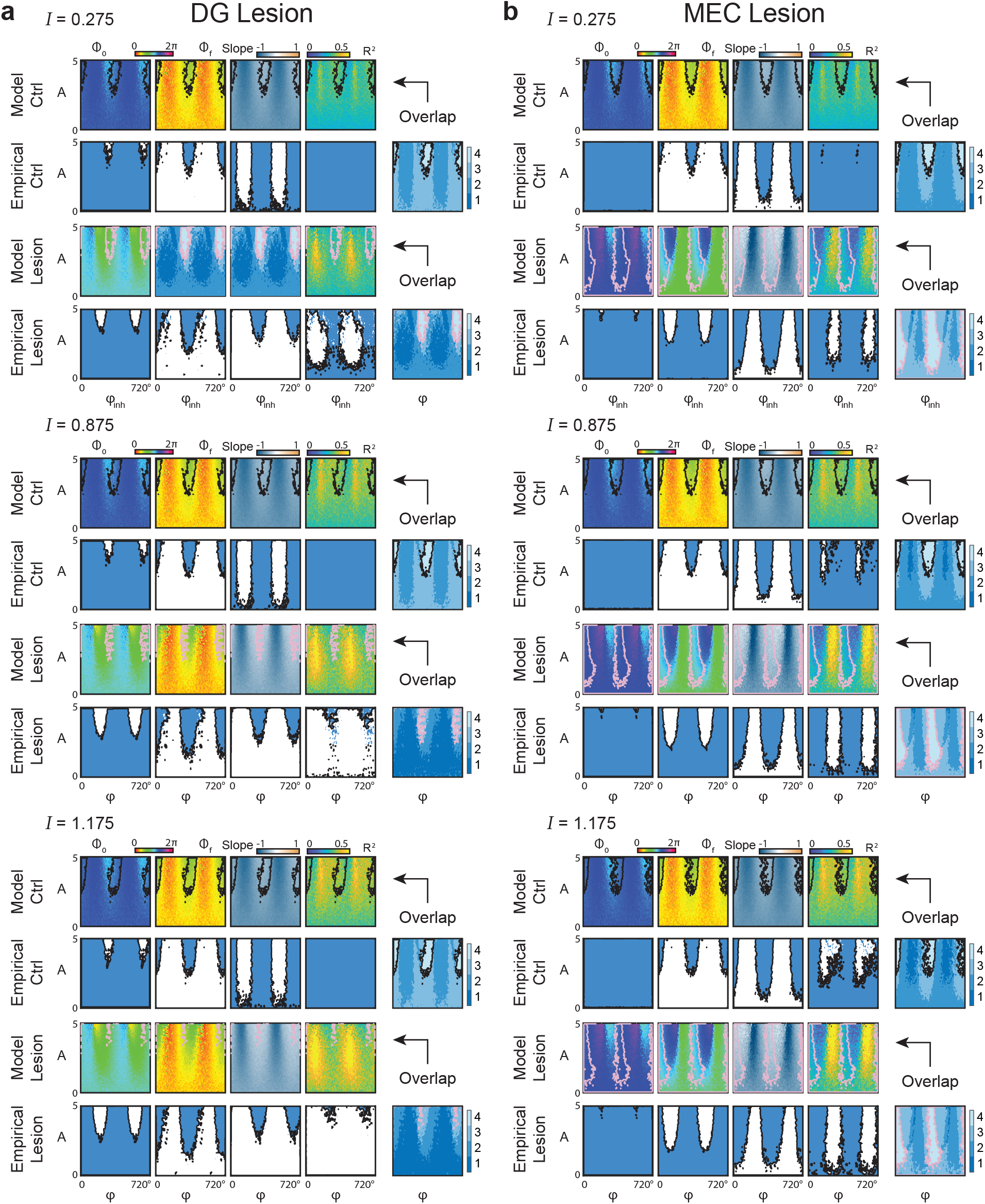
Baseline inhibition level (I) does not qualitatively alter the model state space. Model state space was explored by linear search along the I parameter. Three instances for each experiment are shown as described: (**a**) The state space obtained at I = 0.275, 0.875, or 1.175 was compared to empirical CA3 data in DG-lesioned and associated control rats. **b**, as in **a** but for empirical CA3 data in MEC-lesioned and associated control rats. The overlap plots across all panels reveal matches of the state space for parameters A and ϕ that resemble those reported in Fig. 7, indicating minimal effect of the baseline inhibition parameter (I).

**Table S1.** Summary of rats used in this study.

| Rat ID | Experiment | Experimental Condition | # Qualifying CA3 Units | # Qualifying DG Units | # Sessions |
| --- | --- | --- | --- | --- | --- |
| 122 | DG | CTRL | N/A | 5 | 1 |
| 147 | DG | CTRL | N/A | 6 | 1 |
| 194 | DG | CTRL | N/A | 4 | 1 |
| 600 | DG | CTRL | 50 | N/A | 2 |
| 601 | DG | CTRL | 19 | 2 | 2 |
| 632 | DG | CTRL | 7 | 1 | 2 |
| 650 | DG | CTRL | 8 | N/A | 1 |
| Total |  |  | 84 | 18 | 10 |
| 599 | DG | LESION | 4 | N/A | 1 |
| 633 | DG | LESION | 5 | N/A | 2 |
| 648 | DG | LESION | 11 | N/A | 2 |
| 649 | DG | LESION | 4 | N/A | 2 |
| 656 | DG | LESION | 2 | N/A | 1 |
| 669 | DG | LESION | 20 | N/A | 2 |
| 672 | DG | LESION | 7 | N/A | 3 |
| 675 | DG | LESION | 7 | N/A | 1 |
| 684 | DG | LESION | 8 | N/A | 2 |
| Total |  |  | 68 | N/A | 16 |
| 3661 | MEC | CTRL | 23 | N/A | 5 |
| 3839 | MEC | CTRL | 11 | N/A | 3 |
| 3840 | MEC | CTRL | 2 | N/A | 1 |
| 3906 | MEC | CTRL | 13 | N/A | 2 |
| 3931 | MEC | CTRL | 4 | N/A | 1 |
| 3958 | MEC | CTRL | 39 | N/A | 3 |
| 3959 | MEC | CTRL | 9 | N/A | 3 |
| Total |  |  | 101 | N/A | 18 |
| 3656 | MEC | LESION | 5 | N/A | 1 |
| 3754 | MEC | LESION | 17 | N/A | 5 |
| 3756 | MEC | LESION | 10 | N/A | 2 |
| 3837 | MEC | LESION | 6 | N/A | 2 |
| 3903 | MEC | LESION | 9 | N/A | 2 |
| 3928 | MEC | LESION | 29 | N/A | 3 |
| 3978 | MEC | LESION | 48 | N/A | 3 |
| 3979 | MEC | LESION | 34 | N/A | 2 |
| Total |  |  | 158 | N/A | 20 |

**Table S2.** Statistics from Wilcoxon rank sum tests performed for Fig. 4d.

CTRL<sup>(DG)</sup> vs LESION<sup>(DG)</sup>:
| BIN | 1 | 2 | 3 | 4 | 5 | 6 | 7 | 8 | 9 | 10 |
| --- | --- | --- | --- | --- | --- | --- | --- | --- | --- | --- |
| **Z-STATISTIC** | -2.79 | -2.78 | -2.83 | -2.75 | -2.98 | -0.79 | -0.19 | -0.42 | -1.36 | -1.02 |
| **RANK SUM** | 4863.5 | 3906 | 4369.5 | 4113 | 4695.5 | 4916.5 | 5374.5 | 5353.5 | 4825.5 | 5006 |
| **P-VALUE** | 0.0051 | 0.0053 | 0.0045 | 0.0058 | 0.0028 | 0.42 | 0.84 | 0.67 | 0.17 | 0.30 |
| **HOLM-BONFERRONI CORRECTION** | * | * | * | * | * | n.s. | n.s. | n.s. | n.s. | n.s. |

CTRL<sup>(MEC)</sup> vs LESION<sup>(MEC)</sup> :
| BIN | 1 | 2 | 3 | 4 | 5 | 6 | 7 | 8 | 9 | 10 |
| --- | --- | --- | --- | --- | --- | --- | --- | --- | --- | --- |
| **Z-STATISTIC** | -2.35 | -0.50 | -1.54 | -1.01 | -0.73 | 0.29 | -0.70 | -1.62 | -0.88 | -0.85 |
| **RANK SUM** | 11024 | 10570 | 10708 | 10138 | 11089 | 11773 | 11314 | 10800 | 10446 | 11557 |
| **P-VALUE** | 0.0183 | 0.6145 | 0.1214 | 0.3105 | 0.4628 | 0.7677 | 0.4791 | 0.1052 | 0.3737 | 0.39 |
| **HOLM-BONFERRONI CORRECTION** | n.s. | n.s. | n.s. | n.s. | n.s. | n.s. | n.s. | n.s. | n.s. | n.s. |

\*, significant at alpha = 0.05
n.s., not significant at alpha = 0.05

